# A continuous viral vaccine biomanufacturing platform utilizing multiple bioreactor configurations

**DOI:** 10.64898/2026.01.09.698222

**Authors:** Justin Sargunas, Bradley Priem, Dylan Carman, Taravat Sarvari, Natalie M. Nold, Vaishali Sharma, Andrew Pekosz, Caryn L. Heldt, Michael Betenbaugh

## Abstract

Scalable, continuous biomanufacturing processes have grown in importance to meet demand for smaller bioreactor sizes, lowered production costs, and improved quality attributes. The Sf9/recombinant baculovirus (rBV) expression system demonstrates promise for virus-like particle (VLP) vaccine and gene therapy production. Here, we present a continuous rBV platform integrating an infection plug flow reactor (PFR) between stirred tank growth (gCSTR) and production (pCSTR) bioreactors. Cell expansion in the gCSTR included a ramp-up stage followed by continuous growth, reaching a steady state of 5×10^6^ cells/mL and >90% viability. Péclet number-fit tracer studies confirmed near-ideal plug flow in the PFR, yielding a 10 h residence time and progressive infection as measured by gp64 signaling. Finally, a pCSTR with a residence time of 48 h exhibited sustained recombinant protein production. An integrated pilot cascade incorporating all reactors ran continuously for 5 days, maintaining stable CSTR cell densities and a measurable increase in infected cell diameter from 14.5 μm to 16.1 μm. Western blotting and EM of ∼100 nm VLPs in pCSTR effluent demonstrated platform success. Digital twin mechanistic models across four distinct stages of bioreactor operation and Hill-type relationships for rBV infection kinetics predicted cell growth and death for a 7-day run, demonstrating promise for designing continuous systems in silico and building a quantitative framework for scale-up and optimization. Our multi-stage reactor configuration represents a cell host- and product-agnostic production scheme, particularly for processes prone to product heterogeneity, and paves the way towards a true end-to-end continuous platform for myriad modalities in the future.

## 1. Introduction

Influenza A (IAV) is a virus from the Orthomyxoviridae family that is primarily responsible for both global annual influenza outbreaks and pandemic events due to its ability to mutate and avoid pre-existing immunity [1], [2]. Affecting 5-30% of the world’s population annually, IAV is a public health threat that must be continuously addressed [3]. Vaccines against IAV are essential in limiting disease occurrence and severity but the traditional, and most commonly used, manufacturing process involves injecting embryonated chicken eggs with live IAV virus. Live virus is harvested, inactivated, partially purified, and used as a vaccine in humans [4]. Developed in the 1940s, this egg-based production format has been effective in combatting annual influenza outbreaks. However, this approach possesses several drawbacks including amino acid mutations in key viral proteins due to egg adaptation and protein deformation during chemical inactivation [5], [6], [7]. These limitations lead to reduced vaccine efficacy, especially for immunocompromised and elderly subpopulations [8].

A further, key drawback for current egg-based production processes of IAV vaccines is a lengthy-development time. Averaging around 6 months from construct identification to vial filling, production processes rely on expert predictions of IAV variants that are expected to be dominant in the upcoming influenza season [9]. While these predictions are often accurate, the highly mutative nature of hemagglutinin (HA), the primary IAV antigen involved in human immune response, results in the possibility of misalignment between the vaccine and circulating virus HA sequences [2]. To minimize the risk of vaccine mismatch, production formats with shorter development times that can be adapted with the most up-to-date HA variant must be developed.

Cell-based vaccine production platforms have emerged and gained popularity as alternatives to manufacturing in eggs. One such cell-based process is the *Spodoptera frugiperda* (Sf9)/recombinant baculovirus (rBV) expression system, which has been used to generate FDA-approved vaccines. These include free antigen, nanoparticle, and virus-like particle (VLP) vaccines such as FluBlok (IAV), Cervavix (HPV) and Nuvaxovid (COVID-19) [10], [11], [12]. This cell-culture based format relies on high levels of recombinant protein expression achieved through rBV viral infection of Sf9 cells. In the context of IAV VLP production in Sf9 cells, rBV has previously been engineered to drive the expression of HA and matrix protein 1 (M1) [13]. Hemagglutinin acts as the antigen needed to generate protective immune responses and M1 provides the essential physical internal structure for the IAV [14], [15], [16]. When expressed in the Sf9/rBV system, these recombinant proteins self-assemble into VLPs and function as a replication-deficient, self-adjuvating vaccine by displaying HA to promote the generation of neutralizing antibodies [17]. The development time from identification of an HA variant sequence to injection into patients for an Sf9/rBV vaccine is nearly half that of traditional egg-based production methods [18]. This improvement in development time allows for IAV vaccines generated in Sf9 cells to better account for the antigenic drift of HA, a key driver of vaccine inefficacy in traditional egg-based formats. This faster turnaround time further allows for a highly adaptable cell culture platform to generate IAV vaccines that are more related to current HA variants circulating in the public.

To further shorten production development times of an IAV vaccine, Sf9/rBV manufacturing platforms are amenable to transition into continuous formats [19]. The switch to continuous bioprocessing is of significant interest across the biopharmaceutical and vaccine industries, as this culture mode presents key advantages including: (i) reduced plant footprint, (ii) streamlined upstream and downstream processing to shorten production times, (iii) increased process quality understanding and control, and (iv) reduced capital and operating costs as compared to traditional batch mode processes [20]. The development of continuous cell culture processes for IAV VLP production is particularly promising given recent advances in continuous downstream virus purification systems, such as continuous chromatography [21] and aqueous two-phase systems (ATPS) [22], [23], [24] to purify a range of viruses. Integrating a continuous rBV/Sf9 cell culture platform with such a downstream process could result in a fully continuous, end-to-end biomanufacturing process of an IAV vaccine that can be highly adaptable to current HA variants. This process has the potential to establish a vaccine platform to greatly decrease development time [25], increase space-time yield [26], and more closely match new HA variants [2], as compared to traditional egg-based approaches.

A key challenge in designing a continuous format for Sf9/rBV expression systems arises from the complex nature of rBV infection [27]. Protein production in Sf9 cells relies on infection by an rBV that possesses full sequences of its baculovirus chromosome (bacmid). This full-sequence bacmid encodes for baculovirus proteins needed for viral replication, as well as recombinant genes that yield proteins of interest – in the case of IAV VLPs, these include HA and M1. The recombinant genes, as well as bacmid itself, are replicated utilizing the endogenous Sf9 cell machinery throughout different stages of viral infection. Depending on the production and replication demands within an Sf9 cell, errors in bacmid replication can occur, yielding truncated or recombined sequences that possess incorrect or terminated transgenes. Previous work has shown that these faulty bacmids arise with multiple infection of insect cells with many baculoviruses [28]. New baculoviruses that then package these incorrect target sequences are known as defective interfering particles (DIPs) and have negative impacts on recombinant protein production. This process of DIP generation is known as the Von Magnus phenomenon [29]. Although DIPs themselves are replication-deficient, they can co-infect Sf9 cells alongside intact, functional rBV and compete for cellular resources. Because this results in reduced productivity and increased product heterogeneity [30], DIP formation and competition must be minimized in a continuous process mode.

DIPs pose a considerable challenge in existing continuous Sf9/rBV systems that utilize continuously stirred tank reactors (CSTRs) or perfused systems; these formats create a broad distribution of cell infection times and conditions that lead to a loss of recombinant gene copies and protein titers [19]. As a result, a novel bioreactor format must be utilized that prioritizes primary infection of Sf9 cells with a constant, low multiplicity of infection (MOI) to ensure consistent infection conditions. Prior studies have shown that a plug flow reactor (PFR) can be used to (i) avoid the Von Magnus effect during generation of live influenza virus in Madin-Darby canine kidney (MDCK) cells and AGE1.CR.pIX cells [31] and (ii) infect Sf9 cells for short residence times at high multiplicity of infection [32]. In the latter study, an improvement in batch protein expression following PFR infection was observed [33], demonstrating the potential upside of utilizing this technology. Despite these initial successes, this style of reactor has never been applied in the context of end-to-end continuous manufacturing through integration with other bioreactors or unit operations.

In this work, we integrated a PFR-based continuous rBV infection of Sf9 cells into a novel 3-stage bioreactor format to produce an influenza vaccine in the form of a VLP (**Figure 1**). The key challenges in designing this process arose from the complex interplay between (i) cell growth and health, (ii) viral infection and propagation, and (iii) the subsequent production of recombinant proteins to form VLPs. We addressed each of these challenges through reactor optimizations, and then successfully performed a proof-of-concept production run to generate IAV VLPs. Using development and process data, we developed mechanistic models to describe, then predict future campaigns. Taken together, this work demonstrates the feasibility of a continuous upstream process for the generation of not only influenza VLPs, but a platform agnostic to cell line and biologic product.

**Figure 1:**
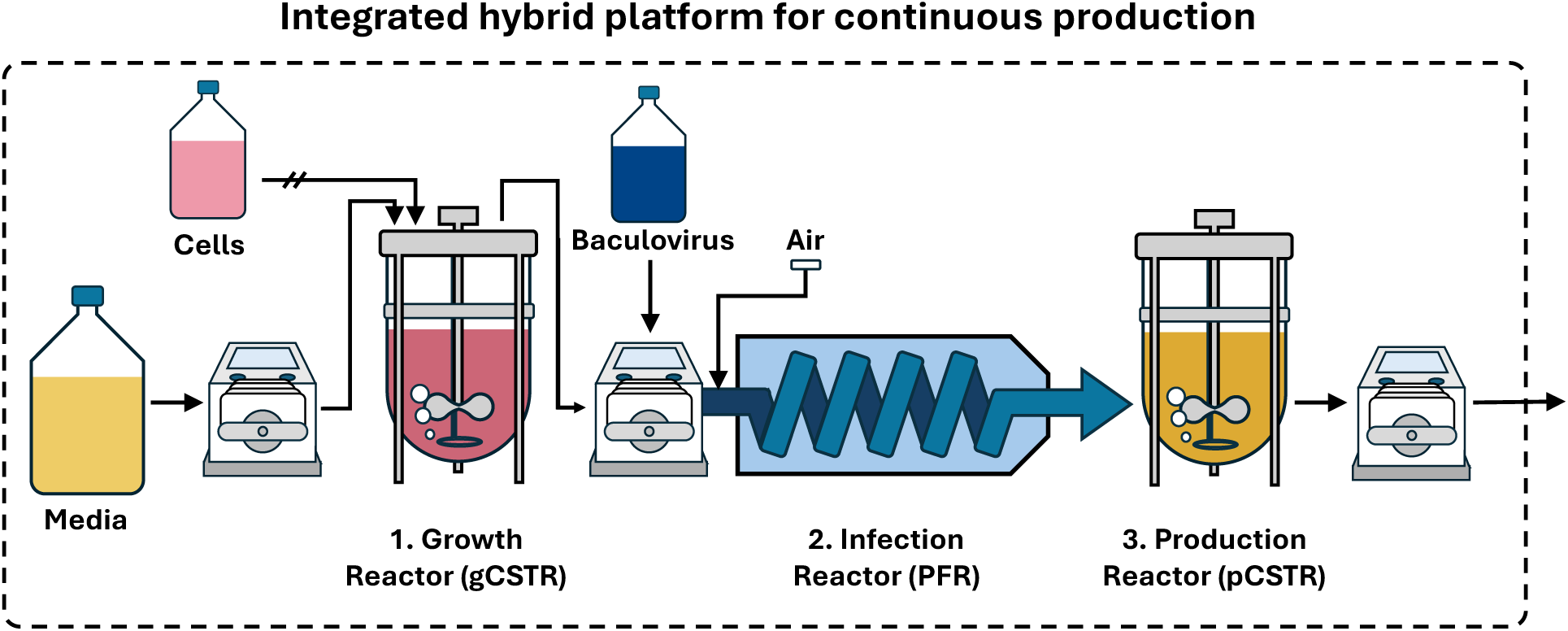
Overview of continuous biomanufacturing platform utilizing the insect cell-baculovirus expression system. Sf9 cells were grown to target density in the growth reactor. Once target density was reached, continuous operation was activated, diluting Sf9 cells from the growth reactor into the infection reactor where recombinant baculovirus was introduced to initiate infection. After Sf9 cells were infected over the length of the infection reactor, they were deposited into the production reactor, in which they produced recombinant protein, with Influenza A VLP generation used as a proof of concept.

## 2. Materials and Methods

### 2.1 Sf9 cell culture

ExpiSf9 cells (Cat No. A35243) and ExpiSF CD medium (Cat. No. A3767802) were obtained from ThermoFisher (Waltham, MA, USA) and cultured as recommended by the manufacturer [34]. In brief, shake flask cultures were inoculated in 125 mL flat base Erlenmeyer flasks (Cat. No. NST-781011) from Diagnocine (Hackensack, NJ, USA) at a target seed viable cell density (VCD) of 0.5×10^6^ cells/mL in a working volume of 30 mL. Sf9 culture was scaled up to 200 mL working volume in 1L shake flasks (Cat. No. NST-784011) from Diagnocine (Hackensack, NJ, USA) to prepare bioreactor seed trains. VCD, viability, and live cell diameter of Sf9 cells were measured with an automated Countess 3 cell counter (Cat. No. A49862) from Invitrogen (Waltham, MA, USA). Cells were cultured in a non-humidified, non-CO_2_ regulated incubator at 27°C away from light in an Innova 4080 incubator shaker (Cat. No. NB-4080) from New Brunswick Scientific (Edison, NJ, USA).

### 2.2 Construction of recombinant baculovirus constructs

Generation of recombinant baculoviruses was performed with the Bac-2-Bac Baculovirus Expression System kit (Cat. No. 10359016) from ThermoFisher (Waltham, MA, USA) per manufacturer instructions. The bicistronic donor plasmid used for bacmid generation, pFastBac-Dual-M2, was a gift from Xiao-Wen Cheng (Addgene plasmid # 135584; http://n2t.net/addgene:135584; RRID: Addgene_135584).

Gene sequences for hemagglutinin and matrix protein 1 were kindly provided by Dr. Andy Pekosz (School of Public Health, Johns Hopkins University) from A/California/7/2009 and A/Udorn/307/1972, respectively. For HA and M1 insertion into pFastBac-Dual-M2, the donor plasmid was linearized using double restriction digest followed by Gibson Assembly with a NEBuilder HiFi DNA assembly master mix (Cat. No. E2621) from New England Biolabs (Ipswich, USA). Gene sequences were amplified with the following primers to yield homologous overhangs to facilitate Gibson assembly cloning (**Table 1**).

**Table 1:**
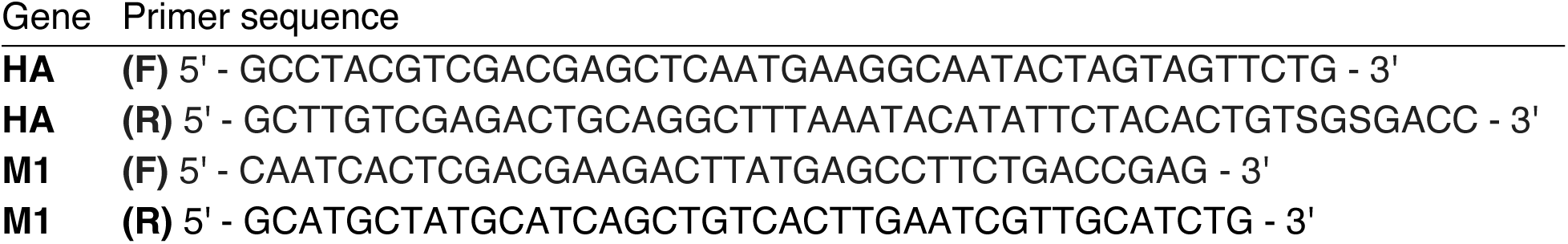
Process variables for mass balance model.

All restriction enzymes were supplied by New England Biosciences (Ipswich, MA, USA). Vector linearization was performed per manufacturer instructions. Gibson assembly was performed as per manufacturer instructions. Confirmation of insertion was performed by via full plasmid sequencing by Plasmidsaurus (San Francisco, CA, USA). A control Gus-containing donor plasmid, pFastBac-Gus, was obtained from ThermoFisher (Waltham, MA, USA). Constructed pFastBac and pFastBac-Dual-M2 donor plasmids were then transformed into Max Efficiency DH10Bac Competent Cells (Cat. No. 10361012) from ThermoFisher (Waltham, MA, USA) following manufacturer protocols. Successful transformants were identified through blue/white screening, in addition to triple antibiotic selection with kanamycin, tetracycline, and genetecin. Kanamycin (Cat. No. K1377), tetracycline (Cat. No. T7660), IPTG (Cat. No. I1284), and X-Gal (Cat. No. 9630-OP) were purchased from Sigma Aldrich (Saint Louis, MO, USA). Genetecin (Cat. No. 15710064) was obtained from Gibco (Waltham, MA, USA). Single colonies were grown and harvested to purify complete bacmid DNA using PureLink HiPure Filter Plasmid Maxiprep Kit (Cat. No. K210007) from ThermoFisher (Waltham, MA, USA).

### 2.3 rBV generation and expansion in Sf9

To generate passage 0 (P0) rBV, Sf9 cells were seeded at 2.5×10^6^ cells/mL in 25 mL of ExpiCD medium in 125 mL shake flasks. Sf9 cells were transfected using Gibco ExpiFectamine Sf Transfection Reagent (Cat. No. A38915) from ThermoFisher (Waltham, MA, USA), as per manufacturer’s protocol [34]. Once transfected cells reached 60-80% viability, virus-containing supernatants were harvested via centrifugation at 300×g for 5 minutes. Viral titer was calculated via flow cytometry. To generate successive viral stocks, Sf9 cells were infected at MOI = 0.1 viruses/cell. Supernatants were harvested when cells fell to 60-80% viability.

### 2.4 Péclet number fitting via video analysis

To determine Péclet numbers, 1.17-, 2.54-, and 3.17-mm inner diameter silicon tubing (Cat. Nos. 8600-0020, 8600-0040, 8600-0030, respectively) were obtained from ThermoFisher (Waltham, MA, USA). Corresponding 3-stop pump tubing (Cat. Nos. MFLX96461-30, MFLX96461-46, MFLX97619-49) was obtained from Avantor (Radnor, PA, USA). 4-Channel Reglo ICC peristaltic pumps (Cat. No. MFLX78001-82) were obtained from Avantor (Radnor, PA, USA).

Equal residence times of tubing were set against a white background, and a camera was used to record video footage at the tubing outlet prior to the peristaltic pump. For each experiment, the tubing was initially primed and filled with water. At experiment start, a blue tracer dye was introduced stepwise at the PFR inlet. The PFR outlet was video recorded until the dye color was visually observed to stabilize. The concentration of the tracer in the fluid at the PFR outlet was measured using image processing tools in Python 3.13.0 (opencv library). Briefly, a constant region of tubing was selected, and the hue was calculated for every frame. The baseline value from the water blank was subtracted, and hues were nondimensionalized by dividing by the final hue, after confirming that the value had stabilized. These data were fit to a function to calculate and export relevant values such as the Péclet number.

### 2.5 Flow cytometry assay for baculovirus titration and Sf9 cell infection

Baculovirus titration was performed using anti-gp64 antibody (Cat. No. 12-6991-80) from ThermoFisher (Saint Louis, MO, USA) per manufacturer’s instructions and previous work [35]. In brief, rBV stock solutions were collected from infected cell culture by centrifuging the culture at 300×g for 5 min and retaining the supernatant. This rBV-containing supernatant was added to a 24-well plate containing Sf9 cells at 1.25×10^6^ cells/mL in 1 mL working volume. Increasing 1:10 dilutions of the rBV stock were prepared in ExpiSf CD medium, and 200 uL of each dilution was added to the well plate. Cells with rBV virus were then incubated for 14-16 hours at 27°C while shaking at 125 RPM. On day 2, cells were collected, pelleted at 300xg for 5 min via centrifugation, washed with PBS, incubated with APC-conjugated anti-gp64 antibody for 30 minutes, then washed again with PBS and resuspended in a dilution buffer (2% fetal bovine serum in PBS solution). These cells were then analyzed using a BD FACSCanto flow cytometer (Franklin Lakes, NJ, USA) for APC detection (Red laser: Excitation: 633-647 nm; Emission 660 nm). The dilution of rBV stock that presented a percent positive gp64 Sf9 cell number of <10% was then carried forward to quantify rBV titer in infectious viral particles (ivp) per mL:

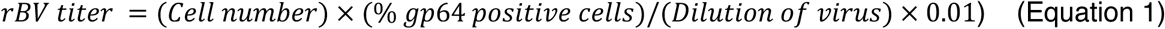

An abbreviated version of this flow cytometry method, beginning at day 2 of the protocol, was used to calculate percent gp64-positive Sf9 cells along the length of the infection PFR to assess infection dynamics in this unit operation. All cytometry data were analyzed using FlowJo 10.8 software.

For fitting of “early” and “late” subpopulations, gp64+ population data were exported and further processed in Python 3.13.0. Intensities were log10-transformed and fit to two Gaussian components via expectation–maximization (scikit-learn GaussianMixture package). For each sample, we computed population percentages and component means (**Supplemental Figure S1**).

### 2.6 β-Glucuronidase (Gus) assay

Extraction buffer and assay buffer were prepared as described in a protocol obtained from Gold Biotechnology (St. Louis, MO, USA) [36]. Sodium phosphate, monobasic monohydrate (Cat. No. 424390250) was obtained from Thermo Scientific (Waltham, MA, USA). Sodium phosphate, dibasic (Cat. No. 424370025) was obtained from Thermo Scientific (Waltham, MA, USA). Ethylenediaminetetraacetic acid disodium salt dihydrate (Cat. No. E4884) was obtained from Sigma Aldrich (St. Louis, MO, USA). Dithiothreitol (Cat. No. R0861) was obtained from Thermo Scientific (Waltham, MA, USA). N-Lauroylsarcosine sodium salt (Cat. No. 434371000) was obtained from Thermo Scientific (Waltham, MA, USA). Triton X-100 (Cat. No. X100) was obtained from Sigma Aldrich (St. Louis, MO, USA). 4-Methylumbelliferyl beta-D-glucuronide (MUG) (Cat. No. ab146354) was obtained from abcam (Cambridge, UK). 4-Methylumbelliferone (4-MU) (Cat. No. M-499-1) was obtained from Gold Biotechnology (St. Louis, MO, USA).

A β-Glucuronidase (Gus) assay was performed to assess Sf9 cell productivity (**Supplemental Figure S2**). Gus-producing Sf9 cells were harvested, and cells were pelleted via centrifugation at 300×g for 5 minutes. Cell pellets were resuspended in extraction buffer and sonicated on ice using a SFX150 Sonifier with Handheld Converter (Cat. No. SFX150) from Branson Ultrasonics (Brookfield, WI, USA) at 20% power with 1 second on/off pulses for 2 minutes to lyse cell membranes. Sonicated samples were centrifuged at 10,000×g for 5 minutes at 4°C. Sample supernatants and 4-methylumbelliferone standards were then prepared as per a protocol from abcam [37] (Cambridge, UK) and loaded into a 96 well Nunc™ F96 MicroWell™ Black Plate (Cat. No. 237107) from Thermo Scientific (Waltham, MA, USA). The plate was analyzed via a Gemini XPS microplate reader (Cat. No. XPS) from Molecular Devices (San Jose, CA, USA), which was operated according to the Abcam protocol to determine Gus activity [37]. A script in Python 3.13.0 was used to determine the Gus activity from the measured fluorescence over time. Two-way ANOVA analysis with Tukey’s multiple comparisons test was utilized to compare Gus activity across experimental conditions, when applicable.

### 2.7 SDS-Page and Western Blotting

Cell culture samples were collected from each bioreactor at each time point. Samples were spun at 300×g, and supernatants were stored at −80°C before use. When necessary, total protein content was measured via bicinchoninic acid (BCA) assay (Cat. Nos. 23225, 23227 and A65453) from Thermo Scientific (Waltham, MA, USA). For analysis via western blot, samples were loaded either at constant volume or constant protein content (25 μg). Sample preparation, SDS Page, protein transfer, and membrane blocking protocols were carried out as previously described [38]. Primary immunostaining against HA (Cat. No. V-314-511-157) at 1:100 (v/v) and M1 (HB-64; American Type Culture Center) at 1:200 (v/v) was carried out at room temperature for 2 h. Both antibodies were kindly gifted by Dr. Andrew Pekosz. Following primary antibody incubation, membranes were washed 3 x 7 min with 0.1% TBS-T. Secondary immunostaining was conducted for 2 h at room temperature with anti-goat antibody (Cat. No. PI-9500) from Vector Laboratories (Burlingame, CA, USA) or anti-mouse antibody (Cat. No. 1706516) from Bio-Rad (Hercules, CA, USA) at 1:3,000 (v/v). Membranes were again washed 3 x 7 min with 0.1% TBS-T. Blots were visualized using the SuperSignal West Pico Plus Chemiluminescent Substrate Kit (Cat. No. 34580) from Thermo Scientific (Waltham, MA, USA). Images were obtained using Image Lab software from Bio-Rad (Hercules, CA, USA).

### 2.8 Stirred tank reactor setup

2 L stainless steel bench top bioreactors and bioreactor controller/towers (BIOSTAT B-DCU) were obtained from Sartorius Stedim (Göttingen, Germany). Silicon tubing of 1.17-, 2.54-, and 3.17-mm inner diameter (Cat. Nos. 8600-0020, 8600-0040, 8600-0030, respectively) were obtained from ThermoFisher (Saint Louis, USA). 4-Channel Reglo ICC peristaltic pumps (Cat. No. MFLX78001-82), along with corresponding 3-stop pump tubing (Cat. Nos. MFLX96461-30, MFLX96461-46, MFLX97619-49) were obtained from Avantor (Radnor, USA). Dissolved O_2_ probes (Cat. No. 237450) were obtained from Hamilton (Reno, Nevada, USA). pH probes (Cat. No. 52000656) were obtained from Mettler Toledo (Columbus, Ohio, USA). Apart from probes, all headplate inserts were manufactured by Sartorius Stedim. 3-blade segment impellers (Cat. No. BB-8847398) were used for culture agitation. Straight line dip tubes (Cat. No. BB-8807850) were used for cell culture seeding, media addition, and cell culture exit in the growth bioreactor and cell culture entrance and exit in the production bioreactor. Tubing for bioreactor inlets (cell seeding and media addition) was capped using Cole-Parmer® 1/8” male and female leur fittings with respective plug caps (Cat. Nos. UX-12028-51, EW-50109-85, EW-50110-00, EW-50110-36).

Sampling ports were integrated with the exit tubing from the growth and production bioreactors using Masterflex® Y-Connectors (Cat. No. MFLX30614-47) with a 4 in. tubing section capped with male leur lock connections (Cat. No. CLS-1399-F50) for syringe sampling. Both bioreactors were sterilized by autoclave prior to cell culture.

For cell culture in a bioreactor format, Sf9 cells were inoculated at 1×10^6^ cells/mL into a 2 L bench top bioreactor in batch format at 1-2 L working volume. Filtered air was introduced through a ring sparger (Cat. No. UNIVESSEL-00011) from Sartorius Stedim (Göttingen, Germany). Air was sparged at a rate of 100 cubic centimeters per minute (ccm) until day 2 post inoculation, at which a dO_2_ setpoint of 40% was initiated. Agitation was maintained at a constant 200 rpm throughout culture. The growth and production bioreactors were sampled twice daily (3-5 mL) for cell density, viability, and other culture measurements. DO and pH measurements were recorded at each sampling time.

### 2.9 Plug flow reactor setup

A plug flow reactor was constructed for use in a bioreactor format. The PFR was constructed using Nalgene^TM^ Pharma-Grade Silicone Tubing (Cat. No. 8600-0030) from Thermo Scientific (Waltham, MA, USA). PFR length (*L*) requirements were determined using flowrates (*F*) exiting the growth bioreactor and tubing inner diameter (*D*) to achieve the target residence time (*τ*).

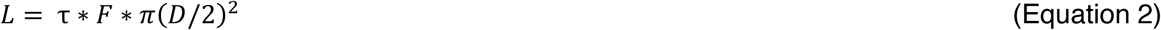

Tubing was generally connected in 25 ft segments, using Masterflex® Y-connectors (Cat. No. MFLX30614-47) to allow for a sampling port at each segment end and feed to the following tubing segment. Three sections of tubing were constructed to allow for 3-stop pump tubing (Cat. Nos. MFLX96461-30, MFLX96461-46, MFLX97619-49) to provide sufficient pump head to maintain consistent flow throughout the PFR. Tubing segments were measured to be 50 ft for segment 1, 75 ft for segment 2, and 50 ft for segment 3. These segments were sterilized using an autoclave, while 3-stop pump tubing was sterilized using 30 minutes of bleach treatment followed by 30-minute washout using sterile water. Once sterilized, these segments and pump tubing were connected using Masterflex® 1/8” Straight Couplers (MFLX40614-50) within a sterile biological safety cabinet. The three stop tubing segments mixing cell culture, baculovirus, and air were also connected to the inlet of PFR segment 1 and were connected to feed bottles containing media and rBV stock in a sterile environment, forming the complete PFR assembly (**Supplemental Figure S3**).

### 2.10 Transmission electron microscopy (TEM)

The morphology of viral particles was characterized by TEM. For sample preparation, a carbon-coated copper grid, NetMesh™ lacey Formvar/carbon stabilized with Carbon, 200 mesh (Cat. No. 01881), from PELCO (Fresno, CA, USA) was plasma-treated (Harrick Plasma, Ithaca, NY, USA) for 15 s at a low RF. The sample was immobilized for two minutes on the grid followed by a one-minute staining with 2% (w/v) uranyl acetate (Cat. No. NC0788109) from Thermo Fisher Scientific (Waltham, MA, USA) prepared in 50% (v/v) ethanol and Nanopure water [39]. Electron micrographs were captured using a FEI 200 kV Titan Themis scanning transmission electron microscope (S-TEM) operated at 200 kV and particle sizing was analyzed using ImageJ software [40].

### 2.11 Kinetic model development

We generated two complementary mechanistic models to describe our continuous system. The first described bioreactor conditions via mass balance, and the second tracked infection progress through the PFR.

For the mass balance model, we defined process variables (**Table 1**), process inputs (**Table 2**), fitted parameters (**Table 3**), and phases of bioreactor operation (**Table 4**).

**Table 1.** Primers for HA and M1 insertion into pFastBac-Dual-M2.

| Variable | Unit | Purpose |
| --- | --- | --- |
| $X_{gCSTR}$ | $10^6$ cells/mL | Viable cell density in gCSTR over time |
| $X_{pCSTR}$ | $10^6$ cells/mL | Viable cell density in pCSTR over time |
| $V_{pCSTR}$ | mL | Reactor volume in pCSTR over time |

**Table 2:** Set process parameters for mass balance model.

| Variable | Unit | Purpose |
| --- | --- | --- |
| $X_{gCSTR,0}$ | $10^6$ cells/mL | Seeding cell density in gCSTR |
| $X_{gCSTR,set}$ | $10^6$ cells/mL | Setpoint cell density in gCSTR |
| $V_{gCSTR}$ | mL | Fixed reactor volume in gCSTR |
| $V_{pCSTR,0}$ | mL | Initial reactor volume in pCSTR |
| $V_{pCSTR,set}$ | mL | Setpoint reactor volume in pCSTR |
| $F_{gCSTR}$ | mL/h | Volumetric flow rate entering and exiting gCSTR |
| $F_{rBV}$ | mL/h | Volumetric flow rate of recombinant baculovirus stock |
| $D_{gCSTR}$ | $h^{-1}$ | Dilution rate of cells from gCSTR |
| $t_{gCSTR,on}$ | h | Time at which continuous gCSTR operation begins |
| $\tau_{PFR}$ | h | Residence time of plug flow reactor |
| $t_{pCSTR,on}$ | h | Time at which continuous pCSTR operation begins |

**Table 3:** Fitted parameters for mass balance model.

| Variable | Unit | Purpose |
| --- | --- | --- |
| $\mu$ | $h^{-1}$ | Specific growth rate for uninfected Sf9 cells |
| $k_{d,inf}$ | $h^{-1}$ | Specific dead rate for infected Sf9 cells |

**Table 4:**
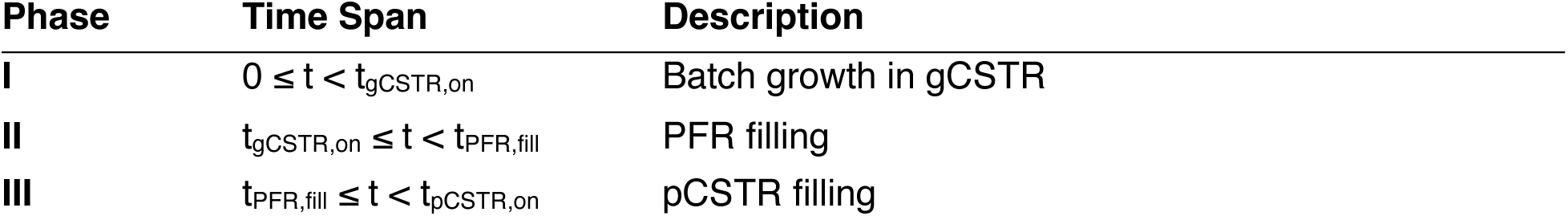
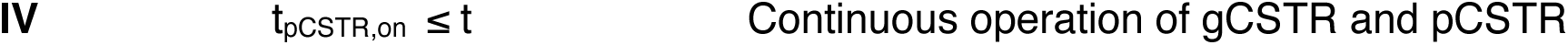
Phases of modelled bioreactor operation.

Process times defining each phase of operation were calculated as:

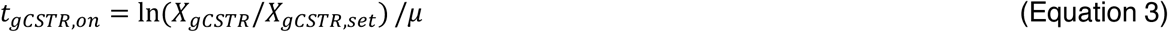

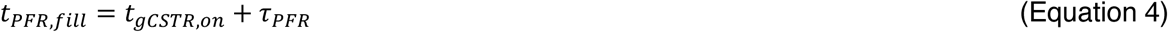

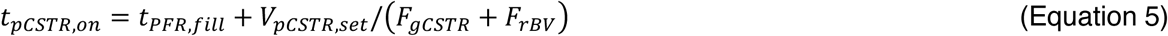

For our mass balance model, we fit gCSTR Sf9 VCD using two piecewise equations, depending on culture mode. For each phase, we assumed a negligible death rate for healthy Sf9 cells. The balance for cells in the initial batch format is:

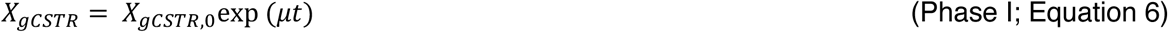

Once the gCSTR reaches a target VCD, continuous CSTR operation commences, and is defined by:

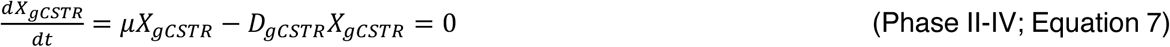

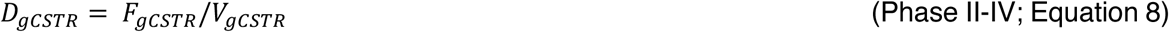

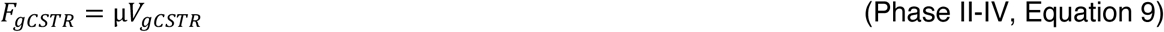

After passing through the PFR, culture enters the pCSTR, which has a volume described by piecewise equations for batch, fed batch, and continuous modes. The volume for each stage is described by:

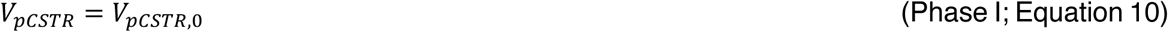

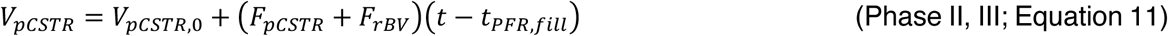

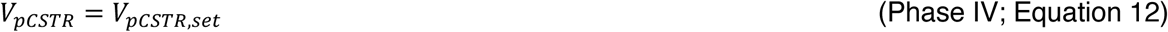

We propose that the cell density exiting the gCSTR is equal to that entering the pCSTR. This arises from the assumption that there is a halt in growth and no retention of cells in the PFR. Thus, biomass in the pCSTR is modelled as:

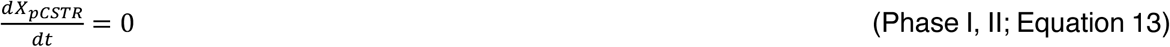

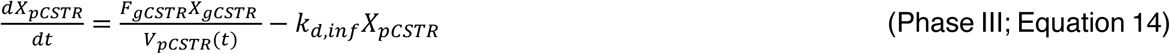

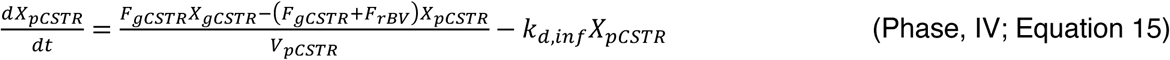

The second mechanistic model describes the infection of Sf9 cells in the infection PFR, with variables fit to a version of the Hill-type sigmoidal equation (**Table 5**) [41].

**Table 5:** Variables for Hill-type sigmoidal fitting of gp64 expression.

| Variable | Unit | Purpose |
| --- | --- | --- |
| $f_{gp64}$ | - | Fraction of Sf9 cells expressing gp64 |
| $f_{max}$ | - | Maximum detectable fraction of Sf9 cells expressing gp64 |
| $k$ | - | Hill coefficient describing viral infection |
| $t_{1/2}$ | h | Time to achieve half of the maximum gp64-positive cells |

We modelled the extent of rBV infection of Sf9 cells as a function of residence time in Equation 16:

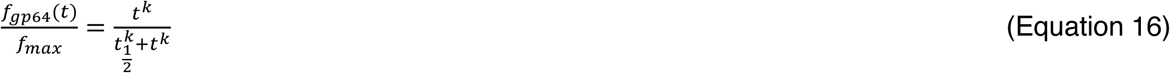

Normalized gp64 assay data from PFR samplings were first nondimensionalized (*f* / *f_max_*). Nonlinear least-squares regression (SciPy curve_fit function) was used to estimate *k* and *t_1/2_*. Parameter uncertainty was assessed by nonparametric bootstrap resampling (n = 1,000). Experimental time-course data were randomly resampled with replacement to generate distributions for *k* and *t_1/2_*, and 2.5^th^ and 97.5^th^ percentiles were used to determine 95% confidence intervals. Model performance we evaluated with an independent validation dataset using the coefficient of determination (R^2^) and root mean square error (RMSE).

## 3. Results

### 3.1 Process overview of 3-stage continuous biomanufacturing platform

Our continuous platform was designed to consist of three stages (**Figure 1**). In the first stage, a stirred tank growth reactor (gCSTR), Sf9 cells were seeded and cultured to a target viable cell density (VCD). Upon initiation of inlet and outlet flows from the gCSTR, Sf9 cells were fed into a tubular plug flow reactor (PFR) for infection. Cells and baculovirus effluent from the PFR were then transferred into a production stirred tank reactor (pCSTR) for extended production. Full integration of the system enabled continuous generation of effluent containing infected cells, baculovirus, and recombinant proteins and IAV VLPs.

### 3.2 Establishing growth reactor conditions for Sf9 cells

A consistent supply of dense and viable cells was a necessary input to our continuous manufacturing system. A stirred tank growth reactor was selected as the first reactor in the cascade to accomplish this task. Cells were seeded in batch format, and after 3 days of exponential growth (t_d_ = 24.8 h), the gCSTR VCD exceeded the target of 5×10^6^ cells/mL (**Figure 2**). From the reactor mass balance (see **Materials and Methods, 2.11**), the dilution rate was set to equal the cell growth rate (*μ* = 0.028 h^-1^), resulting in an initial flow rate of *F* = 0.47 mL/min.

**Figure 2.**
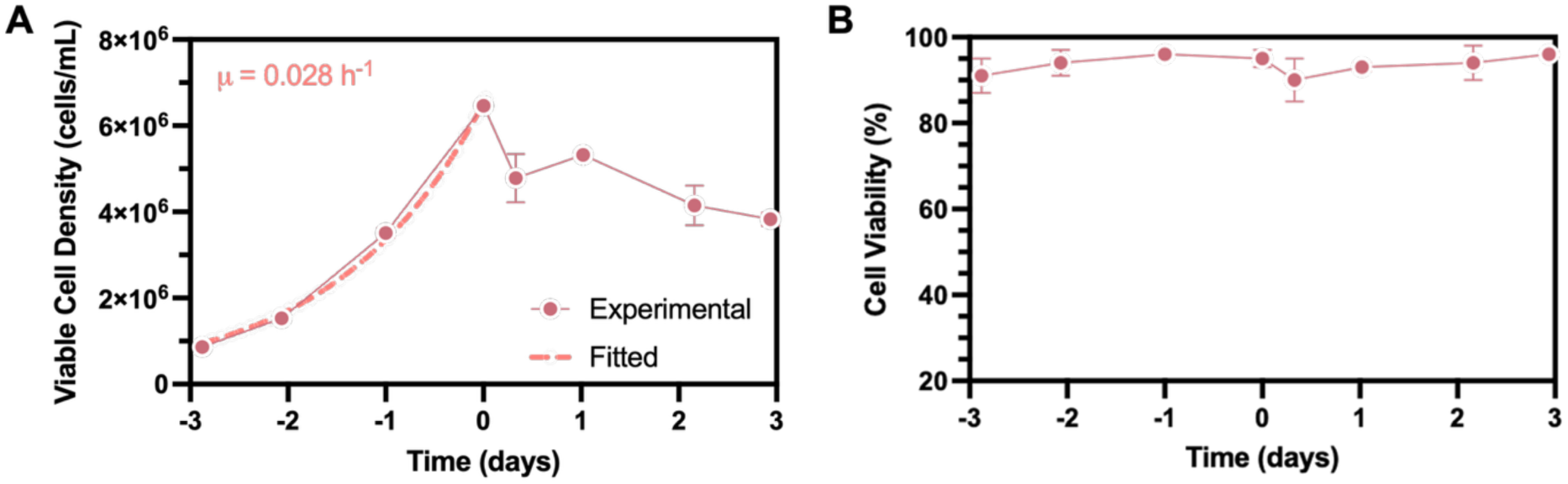
Setup of growth reactor. **(A)** Sf9 cells were cultured in a bioreactor (1 L working volume) for 3 days in batch operation to determine growth rate and achieve a high cell density. Inlet and outlet flows were initiated, and cell density was maintained within 20% for an additional three days, demonstrating success of the CSTR format with manual flow rate adjustment. **(B)** Cells maintained high (>90%) viability throughout both batch and continuous phases of operation (Mean ± SD, n = 2 replicate samplings).

Operating the gCSTR in chemostat mode allowed us to continually remove Sf9 cells at our target VCD while replenishing the lost culture volume with fresh medium, maintaining a target VCD and working volume within the gCSTR during continuous operation. We ran in a continuous mode for three days to determine the feasibility of maintaining the cell culture at a density within 20% of the setpoint (4-6×10^6^ cells/mL; **Figure 2A**). This setpoint was held via manual flow rate adjustment, which ranged from 0.4-0.6 mL/min. High culture viability (>90%; **Figure 2B**) was achieved for the entire 6-day duration of the culture. Dissolved oxygen and pH stayed within expected ranges even at peak cell density, and there was no evidence of depletion of glucose, a potential limiting nutrient, in the gCSTR media at any point (**Supplemental Figure S4**). Having established a continuous feed source of insect cells, we proceeded to investigate ideal conditions for the infection reactor.

### 3.3 Assessing plug flow behavior in infection PFR setup

To support uniform infection kinetics and minimize the productivity loss observed in traditional batch CSTR infection formats, we implemented an infection plug flow reactor (PFR) downstream of the gCSTR to serve as a dedicated rBV infection unit (**Figure 1**). The cell culture flow rate through the PFR was constrained to 0.40-0.60 mL/min, as dictated by day-to-day changes in calculated Sf9 specific growth rate in the gCSTR (**Figure 2A**). To maintain a constant reactor residence time, a total flow rate of 0.75 mL/min was imposed on the system, with rBV and air flow rates adjusted to maintain MOI and total flow rate, respectively.

We began our PFR assessment by comparing the ratio of advective to diffusive flow in different candidates. Ideal plug flow, our desired flow regime, is characterized by fluid that moves in a uniform “plug” dominated by advective forces as opposed to molecular diffusion [42]. This provides consistent rBV and Sf9 conditions at each point along the PFR and facilitates a homogenous infection. To this end, we selected tubing with inner diameters of 1.14 mm, 2.54 mm, and 3.17 mm and experimentally validated the flow profile for each through the fitting of the Péclet number (*Pe*). This dimensionless group compares the relative contribution of advective and diffusive forces as they relate to fluid flow:

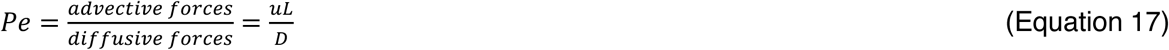

Where *u* is the fluid velocity, *L* is the characteristic length, and *D* is the mass diffusion coefficient. An increasing *Pe* indicates more dominant advection (**Figure 3A**); we selected *Pe* > 100 as the cutoff for ideal plug flow behavior [42]. To quantify flow patterns, we implemented a model system to conduct tracer studies on all three reactor candidates (**Figure 3B**) at a flow rate of 0.75 mL/min, matching the anticipated flow rate of the integrated system. Tracer concentrations were calculated at the PFR outlets (see **Materials and Methods 2.4, 2.9**) and fit to Péclet numbers using a dispersion model [42], [43]:

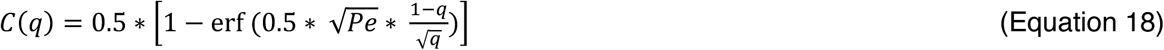

where *C* is the dimensionless concentration and *q* is the dimensionless time. We observed a high *Pe* value across all diameters; at identical volumetric fluid flow rates matching the maximum anticipated gCSTR dilution rate, the tubular reactor with an ID of 1.14 mm had a *Pe* value of 395 (R^2^ = 0.99), the reactor with ID of 2.54 mm had a *Pe* value of 479 (R^2^ = 0.99), and the final with an ID of 3.17 mm had a *Pe* value of 403 (R^2^ = 0.99; **Figure 3C**). We observed a minor divergence between experimental data and the dispersion model at later times (*q* > 1), where the model predicted a sharp, symmetric increase in concentration, while the experimental data showed a “shoulder” with a more gradual increase in concentration. These data implied that the tracer was exiting the PFR at a rate slower than theoretically calculated and suggested that particles in the fluid were retained longer than expected. Previous groups have reported this non-ideality in plug flow in real systems that can result in accumulation within the PFR and back mixing [44]. Nonetheless, we observed high agreement between theoretical concentration changes and the experimental data, and all tubing sizes showed ideal flow behavior as per our definition (*Pe* > 100). While these initial tracer studies were performed in a single-phase liquid setting, subsequent introduction of air segments for cell culture had minimum impact on the flow profile and retained a high degree of axial advective transport, as reactor geometry and flow rate were preserved.

**Figure 3:**
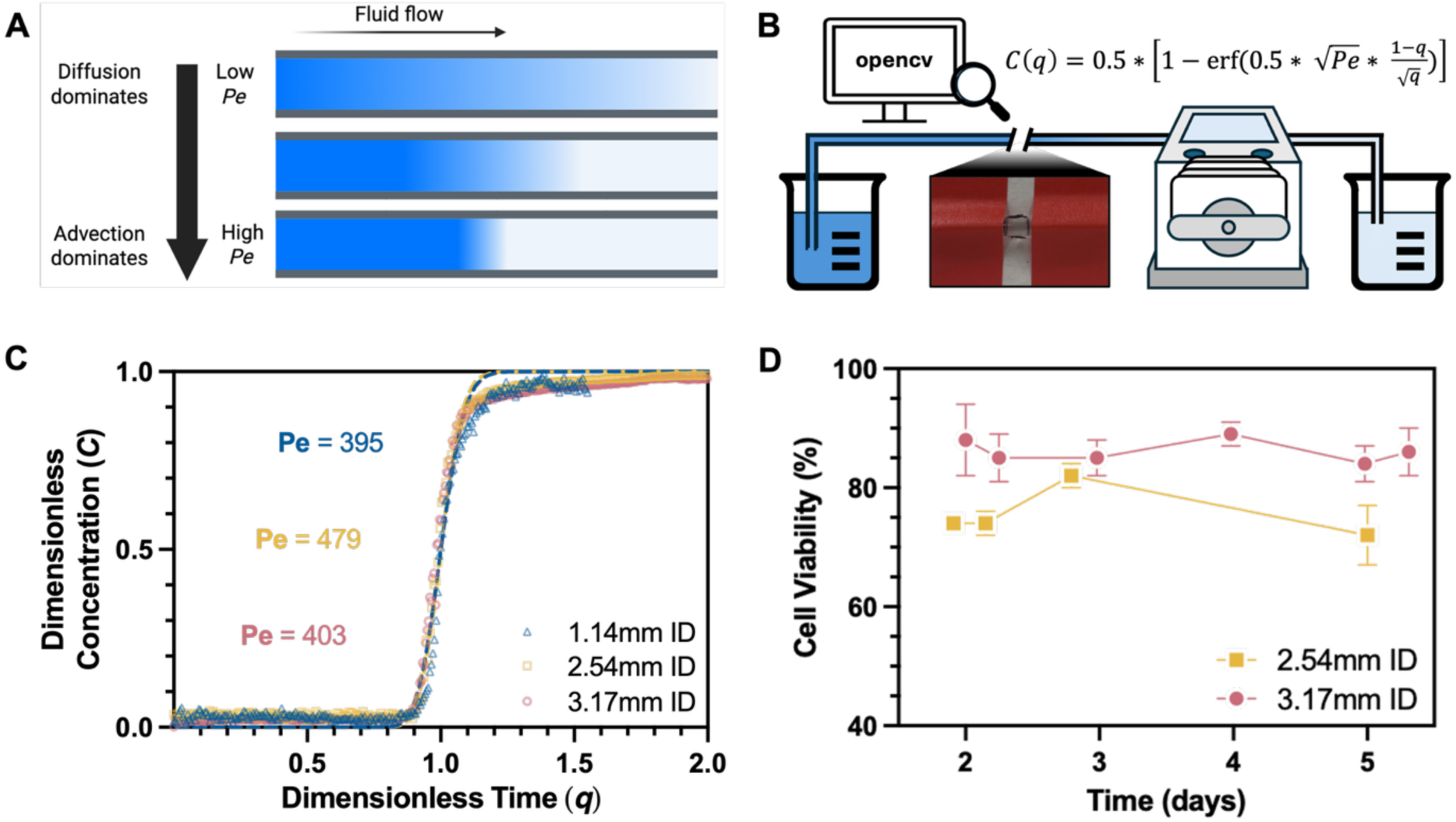
Characterization of plug flow reactor. **(A)** Schematic of different flow behaviors of varying diffusive and advective contributions to fluid flow as described by Péclet (*Pe*) values. **(B)** Experimental workflow of flow studies to determine the ideality of PFR candidates. A dye tracer was introduced stepwise at the reactor inlet and video was recorded at the outlet. Image processing via the opencv library was used to generate time-course concentration data that were then fit to a dispersion model for calculating Pe values. **(C)** Dimensionless concentration data for 1.14-, 2.54-, and 3.17-mm diameter PFR candidates at the reactor exit. Data points show representative experimental data and dashed lines indicate modelled relationship with fitted Pe value for each candidate. **(D)** Cell viability measurements for Sf9 cells following a 10-hour residence time in a 2.54- or 3.17-mm tubular reactor integrated with upstream gCSTR for cell supply (Mean ± SD, n = 2 replicates samplings).

We next evaluated the effect of tubular reactor dimensions on cell viability. For this study, the two larger reactors (2.54 mm and 3.17 mm ID) were carried forward. Sf9 cells cultured in the gCSTR were pumped through either the 2.54mm and 3.17mm ID tubular reactors for a 10-hour residence time, and Sf9 cell viability was assessed for 2-stage integrated bioreactor over a 5-day run (**Figure 3D**). Cells within the 2.54 mm PFR demonstrated consistently lower viability (76% viable, on average) than those in the 3.17 mm reactor (86% viable, on average) perhaps from increased shear stress and hydrodynamic forces [45], [46]. To maintain ideal flow and high cell viability entering the downstream production reactor, we therefore selected the 3.17 mm tubing for our infection PFR.

### 3.4 Identifying ideal infection PFR residence time to maximize Sf9 infection

After selecting the 3.17 mm PFR, we investigated various PFR residence times to characterize the rBV infection of Sf9 cells. We aimed to identify the shortest residence time yielding complete or near-complete infection of Sf9 cells, ensuring cells were infected from fresh rBV stock through a primary infection process, rather than with potentially defective, later-generation viruses in the production reactor.

To assess PFR performance in a cell culture context, Sf9 cells were first cultured to and diluted out of the gCSTR at a density of ∼5×10^6^ cells/mL. Viral infection was accomplished by mixing cells with recombinant baculovirus in line at the PFR inlet, followed by the introduction of sterile air to generate alternating segments of cell culture and gas in order to (i) ensure ample O_2_ supply over the length of the reactor, (ii) limit cell settling [31], and (iii) limit back mixing, as compared to a single phase system [47]. A PFR was constructed that allowed for sampling of intermediate residence times (**Supplemental Figure S3**). Baculovirus was introduced at MOI = 5 viruses/cell [34] in line, followed by sterile air to bring the total flow rate to 0.75 mL/min. After a PFR residence time of up to 10 h, the culture was deposited into a production chemostat. To confirm the mass balance and assess cell health, we measured the VCD and viability of the Sf9 cell culture along the length of the PFR. We observed stable performance in the PFR, with cell densities varying between 4-6×10^6^ cells/mL with an average of 4.7×10^6^ cells/mL (**Figure 4A**). Corresponding gCSTR densities associated with these PFR time points were closely aligned, ranging from 4.7×10^6^ to 5.0×10^6^ cells/mL. This suggested minimal settling or post-infection growth of cells in the tubular reactor. Cell viability in the PFR varied between 81% and 89% with an average of 87% (**Figure 4B**), a small decline from gCSTR viabilities. We next proceeded to quantify the infection kinetics of the reactor.

**Figure 4:**
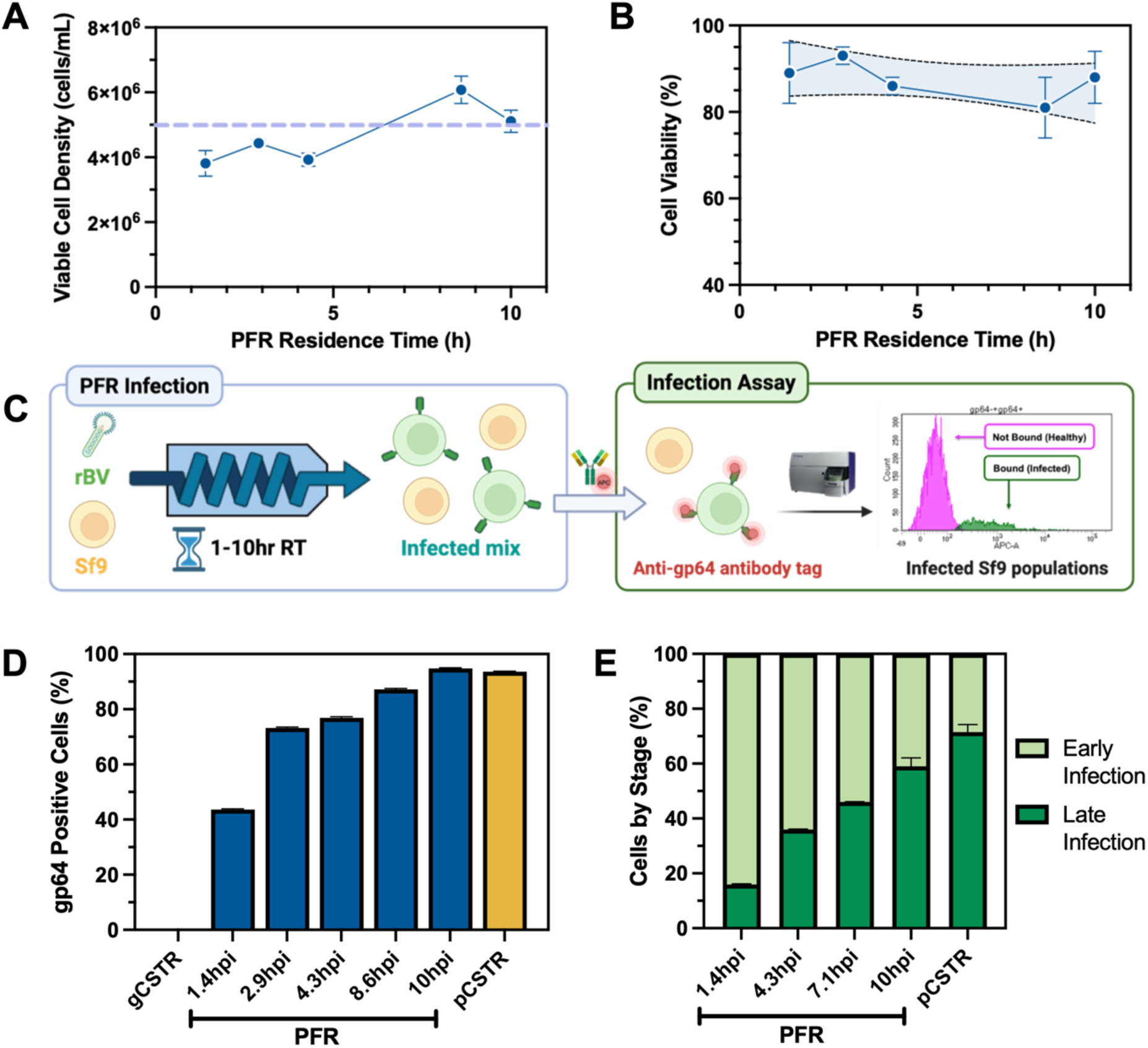
Characterizing PFR performance in a continuous cell cultured format. **(A)** Viable cell density and **(B)** percent viability of Sf9 cells infected with rBV were maintained at ∼5×10^6^ cells/mL and 90%, respectively, for the 10-hour residence time (Mean ± SD, n = 2 replicate samplings). There was no significant trend in viability down the reactor, as determined by linear regression. **(C)** A flow cytometry-based rBV titration method was adapted to determine the progression of cell infection. APC-conjugated anti-gp64 antibody was used to detect infected Sf9 cells displaying gp64 on their surface. APC signal was used to differentiate healthy Sf9 cells (no APC signal) and infected Sf9 cells (with APC signal) to indicate the percent of the total cell population that was infected. **(D)** Application of the flow cytometry assay revealed increasing levels of Sf9 cell infection along the length of the PFR. The percent of infected cells at the PFR outlet matched that of the production reactor (Mean ± SD, n = 2 replicate samplings). **(E)** Analysis of infected Sf9 cell population revealed two subpopulations: early and late infection stages. Increasing levels of late-stage infected cells were observed as residence time in the PFR increased (Mean ± SD, n = 2 replicate samplings). Created with BioRender.

To track infection down the PFR, we adapted a flow cytometric method used for rBV viral titration [35] (see **Materials and Methods, 2.5**) and repurposed it to measure infection kinetics (**Figure 4C**). This assay relies on the surface display of the rBV glycoprotein gp64, an early marker of rBV infection expressed as early as 1-4 hours following initial infection [48], [49]. To measure the extent of infection via gp64 expression, we collected samples of Sf9 cell culture along the length of the plug flow reactor and performed our cytometry method. We observed that the percent of gp64-expressing cells along the PFR increased over time from 43% at 1.4 hpi (hours post infection) to above 93% at 10 hpi on day 5 of continuous operation (**Figure 4D**). Importantly, the 10 hpi gp64+ population of approximately 92% was equivalent to that obtained in the pCSTR, suggesting that the cells had reached the maximum level of infection quantifiable by our assay by 10 hpi. Thus, we chose 10 hours as the residence time of the PFR for our subsequent continuous production experiments.

In this and subsequent PFR runs, we observed two distinct APC-positive Sf9 cell subpopulations **(Supplemental Figure 1**). During early infection PFR residence times, we observed a significant portion of infected Sf9 cells expressing gp64 with a mean APC-A signal of 300 A.U. At later infection PFR residence times, we observed a shift in the infected Sf9 cell subpopulation to a mean APC-A signal of ∼3,000 A.U. These results suggest that Sf9 cells began expressing higher levels of gp64 glycoprotein on their membranes as they entered later stages of infection. Indeed, previous reports suggest that Sf9 cells display increasing extracellular gp64 as their degree of rBV infection increases from early- to late-stage infection [48].

To further investigate these “early” and “late” infected Sf9 cell subpopulations, we fit data to Gaussian curves to estimate the relative abundances of these two phenotypes (see **Materials and Methods 2.5, Supplemental Figure S1**). We assumed that there were only two subpopulation classifications: (1) “early” infection (peak APC-A at ∼300 A.U.) and (2) “late” infection (peak APC-A at ∼3000 A.U.). As infection PFR residence time increased, we observed an increasing abundance of late stage Sf9 cell infection and reducing levels of early stage Sf9 cell infection (**Figure 4E**). At the earliest PFR residence time, 1.4 h, the early Sf9 cell infection population was dominant (84% of all infected cells in early stage). As the PFR residence time increased along the reactor up to 10 h, late infection became the major subpopulation (59% of all infected cells in late stage). Unsurprisingly, the pCSTR itself exhibited an even higher population of late-stage cells (72% of all infected cells in late stage). Together, this analysis shows a clear progression of rBV infection along the PFR and into the pCSTR. Previous work suggests that cells infected by baculovirus significantly downregulate receptors for receptor-mediated viral endocytosis by 24 h, preventing reinfection by progeny virus [50], [51]. A high proportion of late infection cells entering the pCSTR could therefore be advantageous, reducing reinfection by progeny virus and the likelihood of DIP formation.

We note that our expression kinetics differed somewhat from previous reports of surface display of gp64. Namely, we saw significant surface display of gp64 (43% of cells showing expression) and significant late infection populations (16% of infected cells in late stage) as early as 1.4 hpi. These characteristics are typically expected to emerge in the range of 4-8 hpi [48]. This difference likely indicates that some cells were retained along the length of the PFR and experienced a residence time greater than the nominal value based on flow rates. This finding agrees with the behavior observed in the Péclet fitting experiments, in which tracer eluted somewhat slower than expected (**Figure 3C**). Nonetheless, the increasing indicators of rBV infection and consistency in VCD and viability along the reactor indicate that the PFR was a successful tool for controlled infection, and that the retained Sf9 subpopulation was limited.

### 3.5 Confirming pCSTR residence time for optimal protein production

Following growth in the gCSTR and infection in the PFR, the culture mix was transferred to a production CSTR (pCSTR) in which Sf9 cells entered later stages of infection and began producing recombinant protein. Our aim in this stage was to determine an appropriate pCSTR residence time over which infected Sf9 cells could generate recombinant protein while limiting degradation of cellular protein [52].

To assess active protein concentration after infection, we created an rBV construct to drive the expression of β-glucuronidase (Gus) as a model recombinant protein [53]. This reaction was quantified using a fluorescent plate assay, with fluorescent signal intensity being correlated to relative active Gus activity and therefore protein concentration (see **Materials and Methods 2.6**; **Supplemental Figure S2**).

Leveraging this technique, we performed shake flask studies with culture conditions matching our continuous system: Sf9 cells at 5×10^6^ cells/mL were incubated with rBV at MOI = 5 viruses/cell for 10 h, the previously established infection time for the PFR, and 24 h, chosen to probe for the impact of extended infection times on performance. Complete media exchanges were performed after the infection to remove residual baculovirus and emulate the high degree of primary infection seen in the PFR. Cell pellets from this batch format were sampled for productivity via fluorescence assay at 24, 48, 72, and 92 hpi (**Figure 5A**). Elevated Gus activity was observed beginning at 48 hpi for both the 10 h (standard) and 24 h (extended) infection times that persisted through 72hpi (**Figure 5B**). There was no significant difference in Gus activity from 48 to 72 hpi for the standard infection time (p = 0.07), but a minor decrease in activity for the extended infection time for that same period (p = 0.03). Furthermore, Gus activity decreased dramatically at 96 hpi, a 3-fold and 10-fold decrease for the standard (10 h) and extended (24 h) infection time, respectively, indicating a degradation of recombinant protein. Importantly, cultures with extended infection times exhibited lower productivity than the standard time at 48 hpi, 72 hpi, and even 96 hpi (p < 0.0001 for all time points).

**Figure 5:**
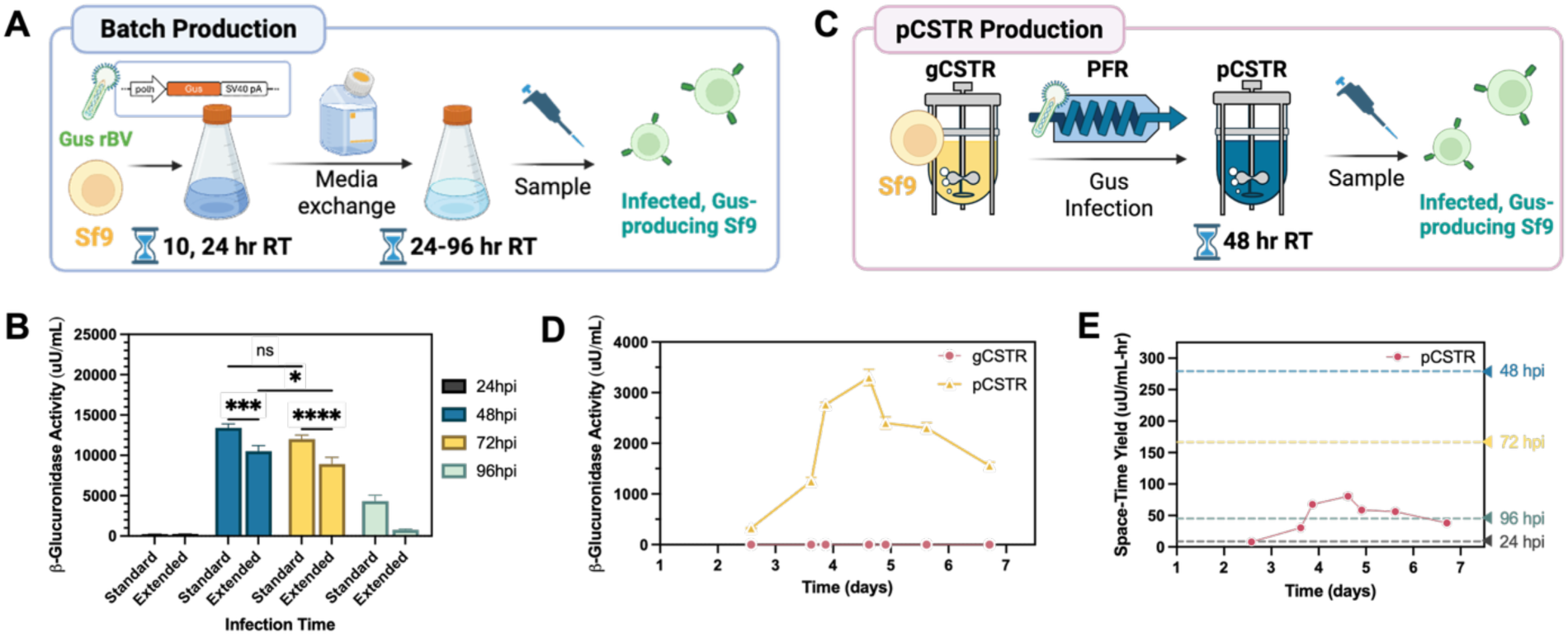
Assessment of recombinant protein production by the measurement of recombinant beta-glucuronidase (Gus) expression in Sf9 cells. **(A)** Batch infection workflow. Sf9 cells were subject to Gus-encoded rBV infection for 10 hours (Standard) or 24 hours (Extended). Following infection, a complete media exchange was performed to remove primary baculovirus. Cell pellets were harvested 24, 48, 72, and 96 hours following initial infection for the Gus activity assay. **(B)** Results from Gus assay on batch infection experiments. Gus activity was highest for standard infection conditions, at 48 and 72 hpi. There was no significant difference in magnitude between these timepoints via 2-way ANOVA with Tukey’s multiple comparisons test (p = 0.07). Extended infection time demonstrated lower productivity at 48 hpi (p = 0.0001) and 72 hpi (p< 0.0001) (Mean ± SD, n = 3 biological replicates; GP: 0.1234 (ns) 0.0332 (*), 0.0021 (**), 0.0002 (***), <0.0001 (****)). **(C)** For continuous operation, Sf9 cells cultured in the gCSTR and infected with Gus-encoded rBV at established PFR conditions (MOI = 5 viruses/cell with 10 hour residence time) were deposited into the pCSTR with a residence time of 48 hours. Following production reactor filling and startup, culture samples were taken from both chemostats over a four-day period for the Gus activity assay. **(D)** Results from Gus assay from continuous chemostat operation. After startup phase, Gus activity levels were consistent over 4 days of chemostat operation in the pCSTR with a target residence time of 48 hours **(E)** pCSTR space-time yield for Gus activity (points and solid lines), as compared to batch (dashed lines) for the process times (Mean ± SD, n = 2 replicate samplings). Created with BioRender.

These batch productivity results supported our choice of the standard 10 h residence time for our infection PFR compared to an extended 24 h residence time, as cultures with extended infection times demonstrated lower productivities at all relevant timepoints tested (**Figure 5B**). Further, these results indicated that 48 h and 72 h incubation times following a 10 h infection were appropriate target residence times for the pCSTR. We selected the 48 h residence time to investigate in our continuous format to reduce the length of viral exposure and reduce the risk of recombinant protein activity reduction from proteases or other mechanisms.

We subsequently applied these parameters to an integrated three-stage bioreactor cascade to assess protein production (**Figure 5C**). Holding conditions in the gCSTR and PFR constant, the pCSTR was set up with a 1.5 L final working volume to achieve a residence time of approximately 48 hours. Following startup and pCSTR filling over the first 2 days of continuous operation, β-Glucuronidase activity peaked at day 4, and remained above 1,500 uU/mL for the remainder of the 7-day run (**Figure 5D**). Converting to space-time yield, the pCSTR achieved a peak productivity of 81 uU/mL-h, as opposed to the range of 45 uU/mL-h (96 hpi) to 279 uU/mL-h (48 hpi) for batch culture (**Figure 5E**). Indeed, while 48 hpi batch condition achieved a higher activity than in the equivalent continuous system in this assessment, a 48-hour residence time was validated as a sufficient pCSTR residence time for protein expression over an extended period of operation [54], [55].

### 3.6 Assessing 3-stage bioreactor format for IAV VLP continuous upstream bioprocessing through pilot run

To demonstrate a pilot run of continuous production of IAV VLPs, we integrated our optimized growth, infection, and production reactors to form a complete 3-stage bioreactor cascade (**Figure 1**). We achieved our target gCSTR VCD of 5×10^6^ cells/mL after 2 days in batch format and maintained that VCD over the next 5 days, the duration of the continuous process (**Figure 6A**). A high cell viability above 90% was maintained in the gCSTR during the entirety of the operation (**Figure 6B**).

**Figure 6:**
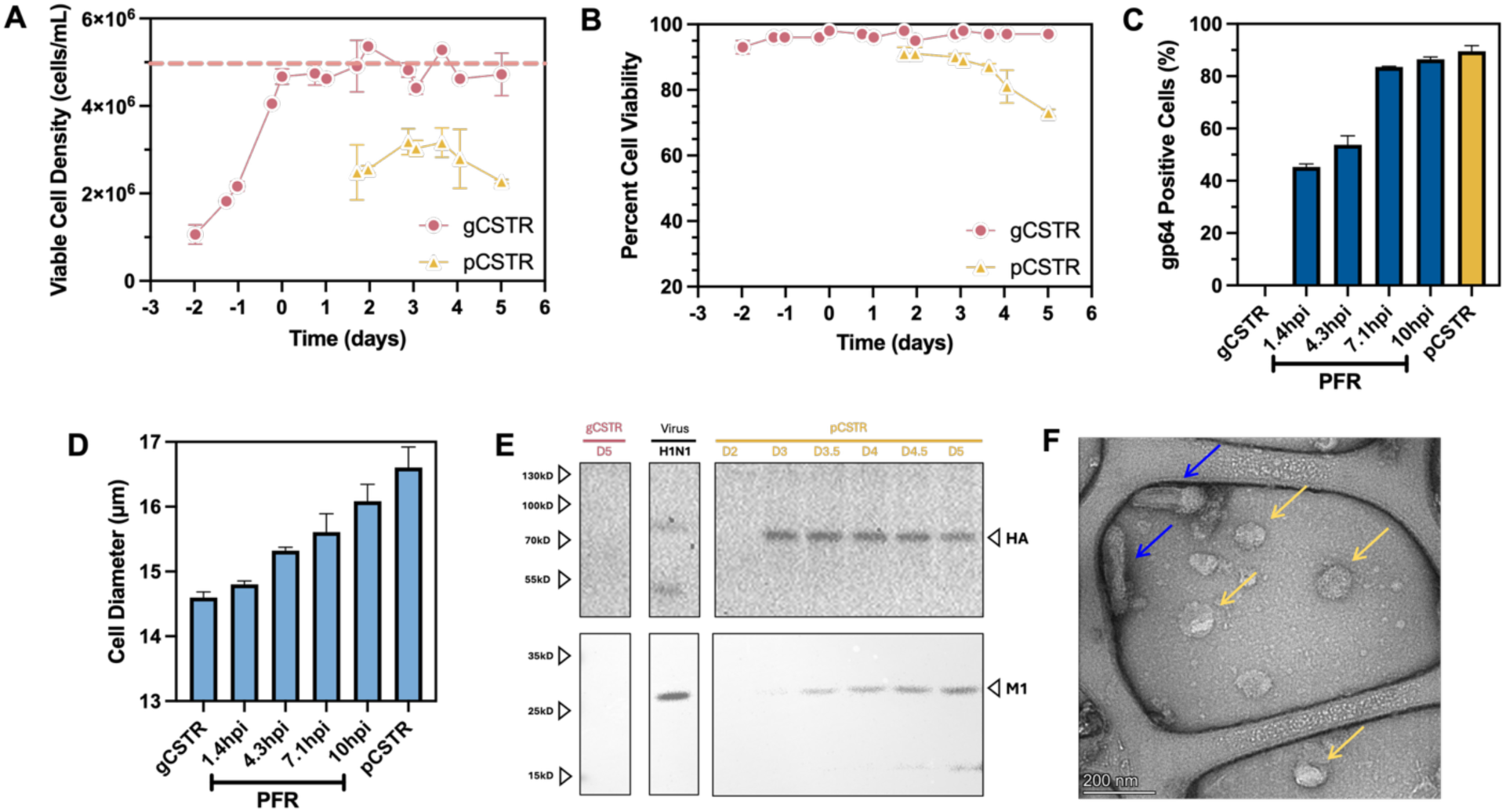
Pilot run of continuous three stage bioreactor system to produce IAV VLPs. **(A)** Viable cell density profiles in the gCSTR and pCSTR remained stable at their expected values after onset of continuous operation. **(B)** A high viability was maintained in the gCSTR and a declining viability in the pCSTR indicated late-stage infection was achieved. **(C)** Sf9 cells samples moving thourgh all three reactors in the cascade showed a shift in cell diameter. Diameters increased along the length of the PFR and into the pCSTR, indicating increasing levels of Sf9 infection. **(D)** Flow cytometry data corroborated an increasingly infected Sf9 cell population along the length of the PFR tubing, consistent with previous data from PFR optimization studies (For all A-D, mean ± SD, n = 2 replicate samplings). **(E)** Western blotting revealed continued expression of recombinant HA and M1 in the pCSTR over the continuous operation. **(F)** Transmission electron microscopy (TEM) imaging suggests the formation of IAV VLPs with an expected diameter of 100 nm (yellow arrows), along with the presence of rBV (blue arrows).

To probe the progression of rBV infection through the PFR, we again measured gp64 binding (**Figures 6C**). As in our preliminary studies, we observed that the gp64-expressing population increased from a moderate size (45%) as early as 1.4 h to a vast majority (85%) at the outlet of the PFR over a 10 h residence time. This gp64+ population was comparable to the 90%-positive population within the pCSTR, suggesting the majority of Sf9 infection occurred through primary infection within the PFR and before the cells entered the production bioreactor.

In addition to gp64 surface display, we measured the Sf9 cell diameter (**Figure 6D**). Increasing Sf9 cell diameter is associated with rBV infection progression and is therefore a useful marker to assess the PFR [56]. Indeed, Sf9 cells displayed a diameter of 14.5 μm in the gCSTR, which increased gradually over the length of the PFR, indicating a progression of rBV infection. At the exit of the PFR, the average cell diameter of 16.1 mm was nearly 10% higher than uninfected cells in the gCSTR, and a 9% increase from the cells at early stages of infection (1.4 h) in the PFR. The average Sf9 cell diameter in pCSTR was even larger at 16.5 μm, suggesting that cells entering the pCSTR continued to expand at late and very late infection periods as a result of viral processing.

Effluent infected Sf9 cells from the PFR entered the pCSTR for extended recombinant protein biosynthesis and VLP formation. After initial transitory operation for filling (days 1 and 2 of continuous operation), viable infected cells were maintained in the pCSTR from days 2 to 5 with densities that exceeded 2×10^6^ cells/mL for the entire continuous run, reaching a maximum of 3×10^6^ cells/mL after day 3 (**Figure 6A**). The lower VCD relative to the gCSTR was attributed to a halt in growth from infection and increased cell death rate in response to viral infection and potential PFR stresses. Given the limitations in process flow rates due to the CSTR mass balance, a 1.5 L pCSTR working volume was selected to achieve a target residence time of approximately 48 hours. Cell viability in the pCSTR declined gradually from 90% to 70% as the Sf9 cells experienced an extended infection period for protein production in the pCSTR (**Figure 6B**). There was a downward trend in viability by days 4 and 5, which likely reflects the challenges of maintaining high cell viabilities of infected cells in a continuous 3-stage cascade.

Finally, Western blotting was performed on pCSTR-harvested supernatants over the course of continuous operation (**Figure 6E**) with HA and M1 expression detected from the pCSTR beginning on day 3. The expression profiles diverged, with rapid peak expression, and then plateau, of HA but gradual accumulation of M1. It is known that membrane and secreted proteins can be difficult to express using a baculovirus system [57]; the greater complexity of hemagglutinin, a transmembrane glycoprotein, could have limited HA expression, while M1, an intracellular cytoplasmic protein, continued to increase.

With confirmed expression of these two Influenza A proteins, we further assessed whether some of these proteins self-assembled into VLPs. To this end, we analyzed ultracentrifuged culture supernatant samples using transmission electron microscopy (TEM) (**Figure 6F**).

Microscopy images showed spherical particles with an average diameter of ∼100 nm (see yellow arrows), matching the expected size of the HA/M1 VLPs [58]. Together, these data demonstrated the expression of HA and M1 as well as the successful formation of IAV VLPs in a continuous 3 - stage bioreactor production process.

### 3.7 Training mechanistic and empirical models of reactor cell growth and infection kinetics

To better characterize our novel bioreactor format, we developed two mechanistic models for cell growth and infection, spanning both CSTRs and the PFR. The cell growth model tracked cell densities and volumes in both stirred tank reactors, with the aim of creating a tool to predict the kinetics to approach steady state under different initial conditions. Cell density data from the pilot run (**Figure 6A**) were used to calculate rate constants: healthy cell growth rate, *μ* = 0.030 h^-1^ and infected cell death rate, *k_d,inf_* = 8.4×10^-3^ h^-1^. Our calculated death rate exceeds those from some previous models [59], [60], which could perhaps be attributed to stresses from travel through the PFR (**Figures 3D, 4B**). These values, together with other process inputs (**Supplemental Table S1**), were used to build the model (see **Materials and Methods, 2.11**).

gCSTR viable cell density (*X_gCSTR_*), pCSTR viable cell density (*X_pCSTR_*), and pCSTR volume (*V_pCSTR_*) were calculated over time from startup of the batch reactor to near-steady state for both reactors using our mechanistic model. Operation was defined in four phases: gCSTR batch operation (I; 2.0 days), PFR filling (II; 0.4 days), pCSTR filling (III; 0.9 days), and fully continuous operation (IV; 3.7 days; **Figure 7A, Supplemental Table S2**). With fitted values from our kinetic expressions (see **Materials and Methods, 2.11**), the overall model (both CSTRs together) showed high agreement with experimental data (R^2^ = 0.97), as well as the values for individual cell densities *X_gCSTR_* (R^2^ = 0.94) and *X_pCSTR_* (R^2^ = 0.97; **Supplemental Table S2**). Steady state cell densities were calculated analytically from the mass balances (**Equations 7, 15**) as 5.0×10^6^ cells/mL and 2.9×10^6^ cells/mL for the gCSTR and pCSTR, respectively. For the pilot run (**Figure 6, Supplemental Table S2**), our model predicted that the pCSTR cell density attained 95% of its steady state value 2.5 days after the onset of continuous gCSTR operation, or 4.7 days total from the beginning of the process. Thus, the modelled bioreactor run showed an approach to, and realization of, a steady state continuous process with respect to cell densities.

**Figure 7:**
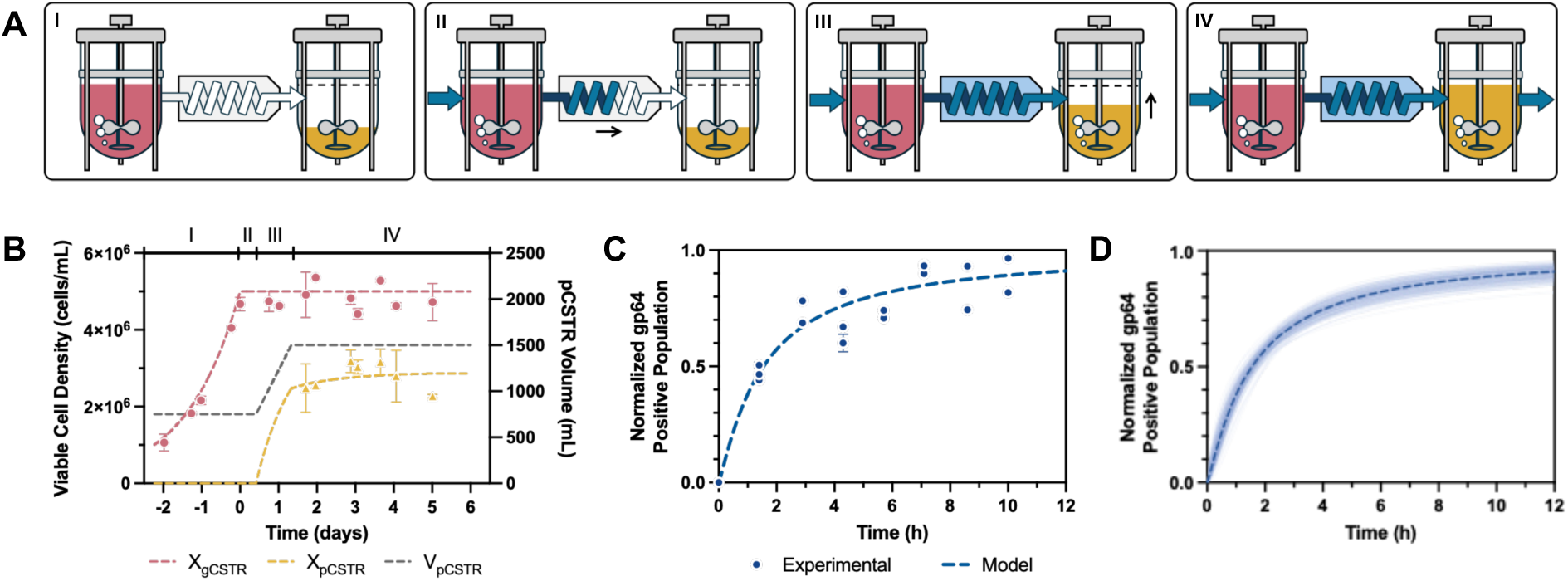
Mechanistic modelling of continuous bioreactor system. **(A)** Four phases of process startup and operation (I-IV): Cells were cultured in batch format in the gCSTR to attain high density (I); after achieving VCD setpoint, inlet and outlet flows were initiated, and the PFR began to fill (II); the partially loaded pCSTR began to fill with material exiting the PFR (III); upon reaching the pCSTR volume target, an outlet flow was initiated, and the process was run fully continuously (IV). **(B)** Predicted viable cell densities in each CSTR (primary axis), and pCSTR volume (secondary axis), as calculated from mass balance for all phases of bioreactor run (I-IV). Model-predicted values (dashed lines) were fitted to experimental data (points) for each phase of operation (Mean ± SD, n = 2 replicate samplings) with high agreement (R^2^ = 0.97). **(C)** Predicted gp64 expression kinetics at MOI = 5 viruses/cell in the plug flow reactor, as fit to a Hill-type equation with data from development and pilot runs (Mean ± SD, n = 2 replicate samplings; data from n = 3 processes) with R^2^ = 0.93. **(D)** Mean predicted Hill relationship from bootstrapped samplings (n = 1,000 subsets) with replacement, plotted with 200 representative fittings from resampling. Parameter error was calculated as 95% confidence intervals from all bootstrapped regressions.

Separately, infection progression was fit to a Hill-type sigmoidal function parameterized by the half-maximal time for gp64 binding (**Equation 16**). Data from three unique data sets – two 10 h PFR development experiments (**Figure 4D**) and the pilot run (**Figure 6D**) – were assessed by nonlinear least squares regression, yielding a Hill coefficient of *k* = 1.1 and time until half response of *t_1/2_* = 1.6 h with R^2^ = 0.93 and RMSE = 0.092 (**Figure 7B**). Indeed, the Hill equation reliably captured infection trends in the PFR and predicted these trends with low error at constant MOI. Due to limited available datasets, we conducted nonparametric bootstrap resampling to assess parameter uncertainty. From 1,000 resampled subsets of data with replacement, we recalculated coefficients of *k* = 1.1 ± 0.3 and *t_1/2_* = 1.5 ± 0.3 h (**Figure 7C**). The value of the Hill coefficient indicates that the gp64 expression used as a proxy for rBV infection followed first order kinetics. Further, there was no cooperative behavior at the MOI and time scales tested, which agrees with past studies showing that an ongoing baculovirus infection does not influence future infections until at least 24 hpi [50], [51]. This span of time is well beyond the tested 10 h PFR residence time and further shows that cells and baculovirus were not significantly retained in the PFR as indicated in our tracer studies (**Figure 3C**). Furthermore, the fitted *t_1/2_* value confirms that infection and gp64 expression occurred rapidly in the PFR, with 50% of cells showing markers of rBV infection as early as 1.5 h after entering the reactor. As a result, these mechanistic and empirical models were able to describe our initial continuous bioproduction study; we next wanted to test the replicability of our findings on a new run.

### 3.8 Validation of kinetic model through independent bioreactor run

To validate our models and demonstrate prolonged continuous operation of the system, we examined a separate integrated continuous 3-stage bioreactor run (**Figure 8**) with minor perturbations to the pCSTR operating conditions (**Supplemental Table S3**). As with previous runs, we were able to maintain a VCD in the gCSTR of ∼5×10^6^ cells/mL (**Figure 8A**) with cell viability >90% (**Figure 8B**). In this case, the pCSTR VCD increased up until day 3 of continuous operation and then remained between 2.2 and 2.5×10^6^ cells/mL over days 4 to 7 (**Figure 8A**). The pCSTR attained 95% of its steady state cell density (2.8×10^6^ cells/mL) after 5.7 days of continuous operation with viability peaking at 70% between days 2 and 3 before settling between 55% and 60% from days 3 to 7 (**Figure 8B**).

**Figure 8:**
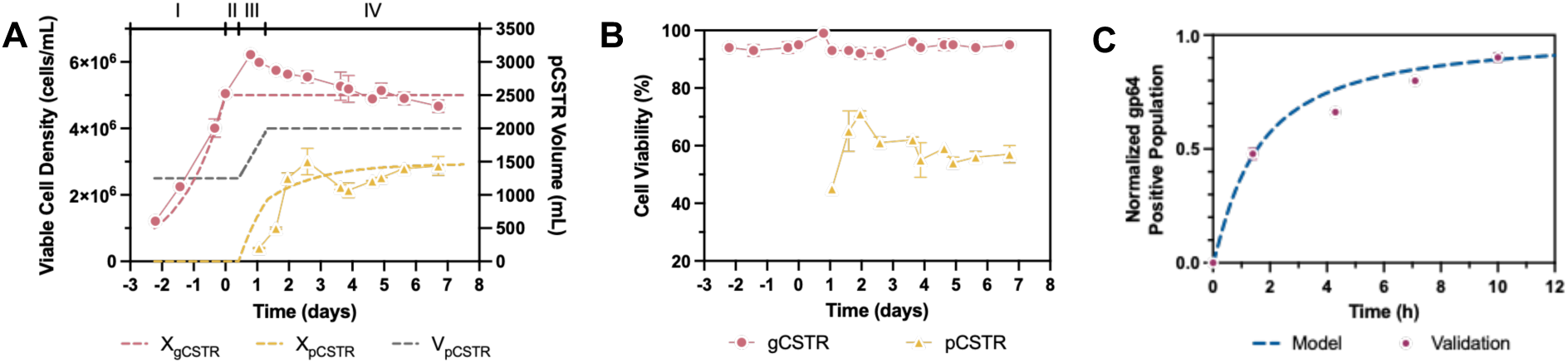
Model performance on validation bioreactor process. **(A)** Predicted viable cell densities in each CSTR (primary axis), and pCSTR volume (secondary axis), as calculated from mass balance for all phases of bioreactor run (I-IV) for validation run. Model-predicted values (dashed lines) were fitted to experimental data (points) for each phase of operation, with good predictive performance (R^2^ = 0. 83). **(B)** Cell viability in the gCSTR exceeded 90% for the entire duration of the process. The pCSTR maintained a viability of 50-60% from days 4 to 7 (Mean ± SD, n = 2 replicate samplings). **(C)** gp64+ population sizes for PFR in validation experiment (points; Mean ± SD, n = 2 replicate samplings) as compared to Hill-type model (dashed line). Predictive performance was exceptional (R^2^ = 0.98).

We next assessed the predictive power of our models on the experimental cell density and infection data for this run (**Figure 8A**). Our cell density model demonstrated good predictive performance for both *X_gCSTR_* (R^2^ = 0.84), *X_pCSTR_* (R^2^ = 0.82), and overall (R^2^ = 0.83; **Supplemental Table S4**). The predicted bioreactor timeline for the four phases of operation (**Figure 7A**) included 2.2 days for batch operation, an additional 1.3 days for PFR priming (II) and pCSTR filling (III), and 5.4 days of operation in the production reactor (IV; **Figure 8A**), comparable to experimental values. We then compared PFR gp64 expression data from the validation run to our fitted Hill-type equation (**Figure 8C**). The model showed strong agreement with the experimental observations and high predictive power (R^2^ = 0.98), with low error (RMSE = 0.05). Together, these metrics confirm that the Hill model is generalizable beyond the training data, and that our parameterized model accurately captures gp64 expression kinetics in the PFR across distinct experiments. These validation data show that we can construct models to mathematically describe all three bioreactors in our continuous system. This may be particularly useful for potential future scaleup considerations, during which cell densities and residence times may shift as system volumes and residence times are perturbed.

To our knowledge, these campaigns and associated models represent the first continuous influenza VLP production process from the Sf9 baculovirus expression system combining 3 integrated stages with separate growth, infection, and production stages. Furthermore, this integrated run marks the longest demonstration of a fully continuous upstream process integrating stirred tank and plug flow reactors via the Sf9 expression system. Our findings, including production of recombinant proteins (**Figures 5D, 5E**) and VLPs (**Figures 6E, 6F**), represent a significant step forward in the path to fully end-to-end continuous Sf9-baculovirus biomanufacturing, which will ultimately include downstream purification of VLPs and other proteins or proteins assemblies.

## 4. Discussion and Concluding Remarks

In this work, we demonstrate a novel, continuous platform for recombinant protein and VLP production with insect cells integrating a plug flow infection reactor between two stirred tank growth and production bioreactors. To drive the continuous expression of HA/M1 VLPs in this growth-infection-production cascade, we expanded Sf9 cells in a gCSTR, uniformly infected them with rBV in a tubular PFR, and continuously generated product in a pCSTR. This was accomplished by optimizing the reactors in stages. Operating the gCSTR as a chemostat supplied a stable stream of Sf9 cells with >90% viability and without nutrient limitations. Our tracer studies revealed that the subsequent infection reactor operated predominantly with ideal plug flow, and a 3.17mm diameter tubing was chosen to maintain high cell viabilities. Using gp64 expression as a surrogate showed that a 10 h residence time at MOI = 5 viruses/cell achieved >90% infection of cells. Finally, a pCSTR residence time of 48 h was revealed from parallel batch studies to balance productivity and sustained protein activity levels. The pilot cascade integrating all 3 reactors ran continuously for 5 days with sustained CSTR cell densities, high infection rates (>85%) and a measurable increase in cell diameter along the PFR. Furthermore, we produced HA/M1 VLPs of ∼100 nm beginning on day 3 lasting until the end of run. Mass balance and Hill equation models, fitted to pilot run data (R² ≥ 0.93), predicted cell growth, death, and infection progression for a subsequent 7-day run, including start-up, transient operation, and approach to steady state. This demonstrated robustness of both the process and the mechanistic framework for scale-up and control for digital twins that can simulate and optimize a complex, multi-unit manufacturing process in silico.

Indeed, this study serves as the foundation for future continuous bioreactor systems for the biomanufacturing of not only influenza but other viral vaccines and cell-based biologics. While we have validated its success in generating VLPs, our reactor configuration represents a cell host-and product-agnostic production scheme, particularly in cell culture processes that are prone to product heterogeneity. While applied here for obtaining consistent infection conditions for rBV and Sf9, this approach can be applied to other time-, shear- and mixing-dependent viral production processes such as generation of viral vaccines in mammalian cell lines like Vero or MDCK as well as transient and stable recombinant adeno-associated virus (rAAV) production. Activation and expansion of lymphocytes or other cell therapies with small molecule inducers is yet another potential application of this platform. These processes see considerable quality variability due to the traditional stirred tank manufacturing format [61], [62], [63], [64] and could benefit greatly from implementation of a well-controlled multi-stage bioreactor system. Our upstream continuous bioreactors can further represent the ideal partners for emerging continuous downstream purification units including continuous chromatography and aqueous two-phase separation (ATPS) systems [21], [23], [24], [65]. In total, this study will pave the way towards a true end-to-end platform for myriad modalities in the coming decades, contributing to efforts in biomanufacturing to streamline processes, reduce production costs, and increase product quality homogeneity for emerging cellular products.

## Data Availability Statement

The data that support the findings of this study are available from the corresponding author upon reasonable request.

## Acknowledgments

The authors would like to thank Dr. Pranay Ladiwala for his assistance and training on bioreactor setup and operation and Dr. Erico Frietas for training on TEM.

## Author Contributions

**Justin Sargunas:** Conceptualization, data curation, formal analysis, investigation, methodology, validation, visualization, writing – original draft, writing – review and editing; **Bradley Priem:** Conceptualization, data curation, formal analysis, investigation, methodology, validation, visualization, writing – original draft, writing – review and editing; **Dylan Carman:** Formal analysis, investigation, methodology, validation, writing – original draft, writing – review and editing**; Taravat Sarvari:** Investigation, methodology; **Natalie Nold:** Conceptualization, investigation, methodology; **Vaishali Sharma:** Investigation; **Andrew Pekosz:** Conceptualization, methodology, resources, supervision**; Caryn Heldt:** Conceptualization, funding acquisition, methodology, project administration, supervision, validation; **Michael Betenbaugh:** Conceptualization, funding acquisition, methodology, project administration, supervision, validation, writing – original draft, writing – review and editing.

---

**Supplemental Figure S1:**
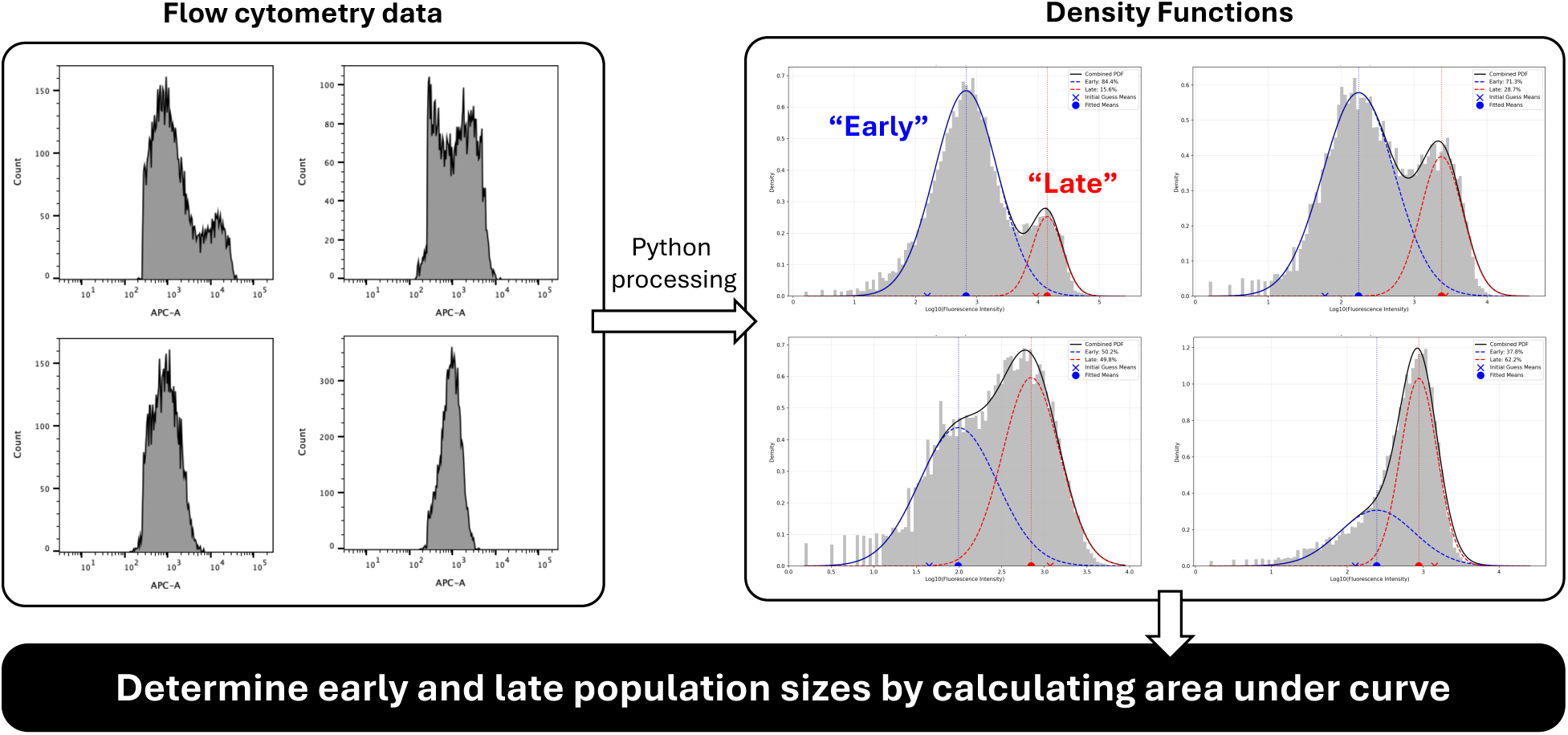
Quantification of “early” and “late” rBV infection. Processed gp64+ flow cytometry data were exported and processed in Python 3.13.0 using the scikit-learn GaussianMixture package. Means and population percentages were calculated for two Gaussian components for each sample.

**Supplemental Figure S2:**
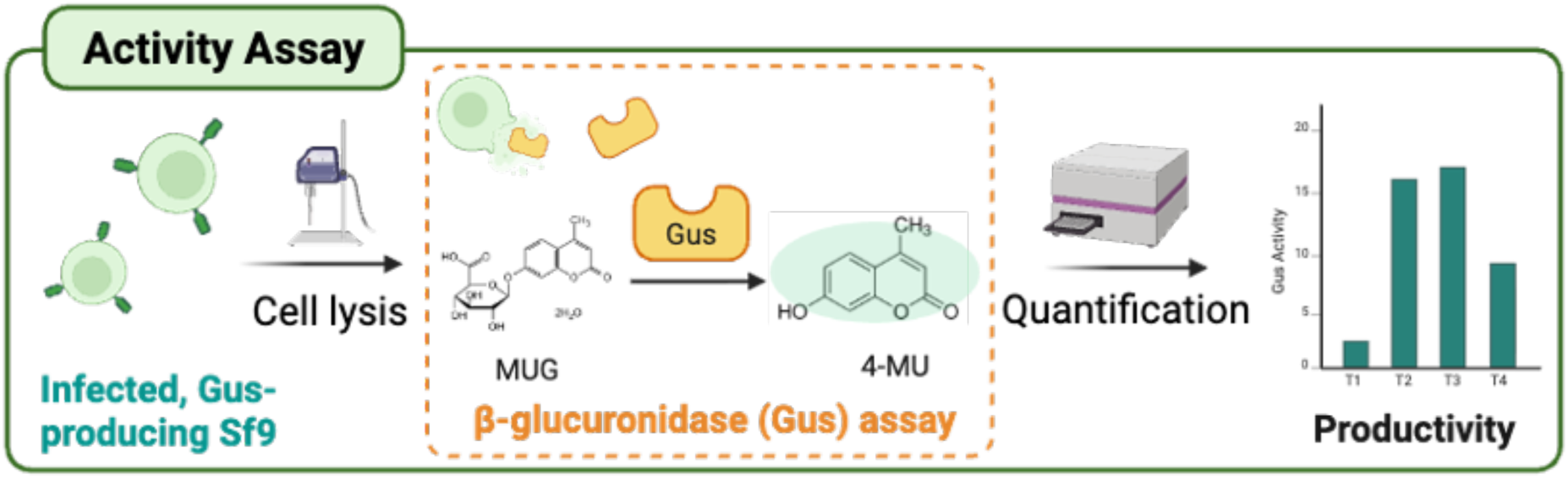
Overview of fluorometric assay. Gus-producing Sf9 cells were lysed via sonication and incubated with MUG substrate to catalyze a reaction to a fluorescent product, 4-MU. Gus activity was calculated from fluorescence increase on a plate assay and correlated to productivity. Created with BioRender.

**Supplemental Figure S3:**
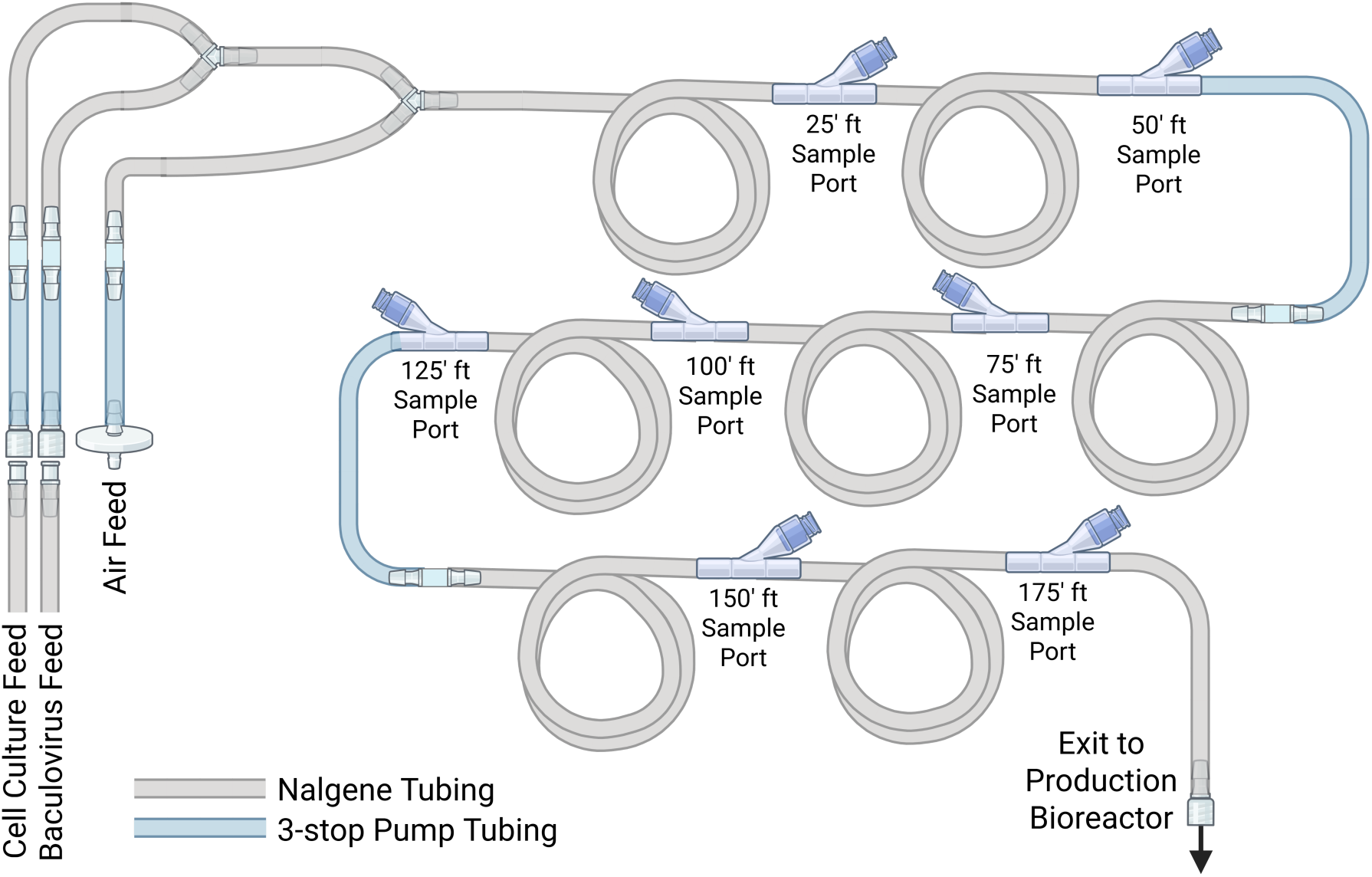
Construction of plug flow reactor. For the 10 h residence PFR, discrete segments of tubing were joined with Y-connectors to allow for sampling along the length of the reactor. To feed the reactor, cell culture was first mixed with recombinant baculovirus, after which air was introduced. Created with BioRender.

**Supplemental Figure S4:**
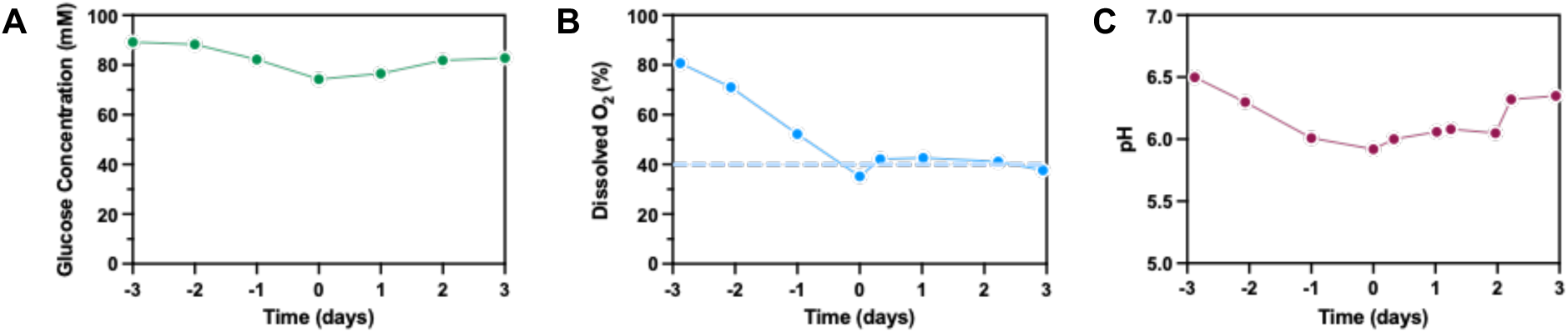
Additional growth reactor characteristics. **(A)** Glucose concentration remained above 70 mM for 6 days, reaching a minimum on day 0 before the start of the continuous operation, but rebounding when flow was initiated. **(B)** Dissolved oxygen content steadily decreased at an aeration rate of 100 ccm from days −3 to −1. After a 40% setpoint was initiated on day −1, oxygen content was maintained through the end of the run on day 3. **(C)** pH of the cell culture similarly declined somewhat during batch operation but recovered when flows began on day 0, returning to a value similar to the start of the run.

**Supplemental Table S1:** Operating values for pilot run and model fitting.

| Variable | Value | Unit |
| --- | --- | --- |
| $X_{gCSTR,0}$ | 1.0 | $10^6$ cells/mL |
| $V_{gCSTR}$ | 1,000 | mL |
| $V_{pCSTR,0}$ | 750 | mL |
| $V_{pCSTR,max}$ | 1,500 | mL |
| $F_{gCSTR}$ | 30.0 | mL/h |
| $F_{rBV}$ | 4.2 | mL/h |
| $D_{gCSTR}$ | 0.030 | $h^{-1}$ |
| $t_{gCSTR,on}$ | 53.6 | h |
| $t_{PFR}$ | 10 | h |
| $t_{pCSTR,on}$ | 85.6 | h |
| $\mu$ | 0.030 | $h^{-1}$ |
| $k_{d,inf}$ | $8.4 \times 10^{-3}$ | $h^{-1}$ |

**Supplemental Table S2:** Model predications versus experimental data for model fitting.

| **Phase** | **Duration (days)** | **gCSTR VCD (E6 cells/mL)** |  | **pCSTR VCD (E6 cells/mL)** |  | **pCSTR Volume (mL)** |  |
| --- | --- | --- | --- | --- | --- | --- | --- |
|  |  | *Experimental* | *Model*<br>*(R<sup>2</sup> = 0.94)* | *Experimental* | *Model*<br>*(R<sup>2</sup> = 0.97)* | *Experimental* | *Model* |
| **I** | 2.0 | 1.1-4.7 | 1.0-5.0 | 0 | 0 | 750 | 750 |
| **II** | 0.4 | 4.7 | 5.0 | 0 | 0 | 750 | 750 |
| **III** | 0.9 | 4.6-4.7 | 5.0 | 0-2.5 | 0-2.5 | 750-1500 | 750-1500 |
| **IV** | 3.7 | 4.6-5.3 | 5.0 | 2.5-3.2 | 2.5-2.9 | 1500 | 1500 |

**Supplemental Table S3:** Operating values for validation bioreactor run.

| Variable | Value | Unit |
| --- | --- | --- |
| $X_{gCSTR,0}$ | 1.0 | $10^6$ cells/mL |
| $V_{gCSTR}$ | 1,000 | mL |
| $V_{pCSTR,0}$ | 1,250 | mL |
| $V_{pCSTR,max}$ | 2,000 | mL |
| $F_{gCSTR}$ | 30.0 | mL/h |
| $F_{rBV}$ | 4.2 | mL/h |
| $D_{gCSTR}$ | 0.030 | $h^{-1}$ |
| $t_{gCSTR,on}$ | 53.6 | h |
| $t_{PFR}$ | 10 | h |
| $t_{pCSTR,on}$ | 85.6 | h |
| $m$ | 0.030 | $h^{-1}$ |
| $k_{d,inf}$ | $8.4 \times 10^{-3}$ | $h^{-1}$ |

**Supplemental Table S4:**
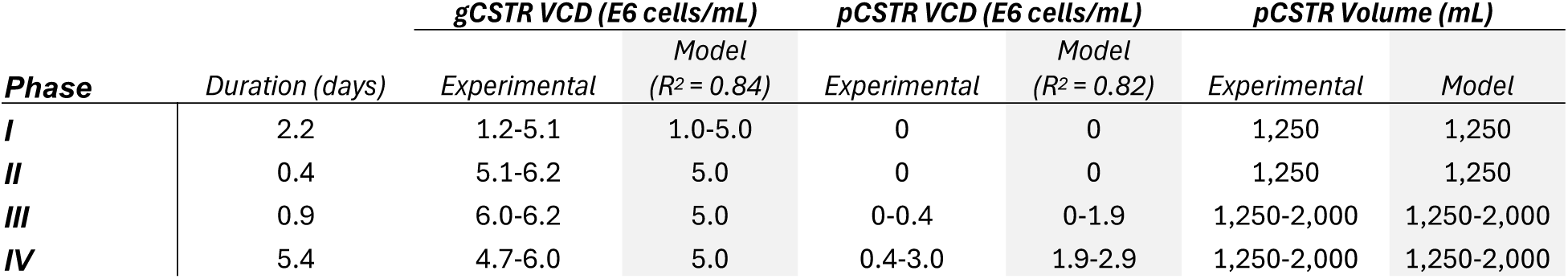
Model predictions versus experimental data for validation run.

## References

[1] D. M. Bell, I. B. Weisfuse, M. Hernandez-Avila, C. Del Rio, X. Bustamante, and G. Rodier, “Pandemic Influenza as 21st Century Urban Public Health Crisis,” Emerg Infect Dis, vol. 15, no. 12, p. 1963, Dec. 2009, doi: 10.3201/EID1512.091232.

[2] F. Carrat and A. Flahault, “Influenza vaccine: The challenge of antigenic drift,” Vaccine, vol. 25, no. 39–40, pp. 6852–6862, Sep. 2007, doi: 10.1016/J.VACCINE.2007.07.027.

[3] T. Horimoto and Y. Kawaoka, “Influenza: lessons from past pandemics, warnings from current incidents,” Nature Reviews Microbiology 2005 3:*8*, vol. 3, no. 8, pp. 591–600, Aug. 2005, doi: 10.1038/nrmicro1208.

[4] D. Gupta and S. Mohan, “Influenza vaccine: a review on current scenario and future prospects,” Journal of Genetic Engineering and Biotechnology, vol. 21, no. 1, pp. 1–10, Dec. 2023, doi: 10.1186/S43141-023-00581-Y/TABLES/1.

[5] C. M. Trombetta, S. Marchi, I. Manini, G. Lazzeri, and E. Montomoli, “Challenges in the development of egg-independent vaccines for influenza,” Expert Rev Vaccines, vol. 18, no. 7, pp. 737–750, Jul. 2019, doi: 10.1080/14760584.2019.1639503;JOURNAL:JOURNAL:IERV20;WGROUP:STRING:PUBLICATION.

[6] W. Liang et al., “Egg-adaptive mutations of human influenza H3N2 virus are contingent on natural evolution,” PLoS Pathog, vol. 18, no. 9, p. e1010875, Sep. 2022, doi: 10.1371/JOURNAL.PPAT.1010875.

[7] J. Herrera-Rodriguez, A. Signorazzi, M. Holtrop, J. de Vries-Idema, and A. Huckriede, “Inactivated or damaged? Comparing the effect of inactivation methods on influenza virions to optimize vaccine production,” Vaccine, vol. 37, no. 12, pp. 1630–1637, Mar. 2019, doi: 10.1016/J.VACCINE.2019.01.086.

[8] A. C. Tricco et al., “Comparing influenza vaccine efficacy against mismatched and matched strains: A systematic review and meta-analysis,” BMC Med, vol. 11, no. 1, pp. 1–19, Jun. 2013, doi: 10.1186/1741-7015-11-153/TABLES/3.

[9] R. M. Anderson, P. J. Scannon, J. T. Matthews, R. Rappuoli, A. Shaw, and D. Estell, “BRIDGE The Planning for Pandemics of Infectious Diseases Pharmaceutical Preparedness for a Pandemic Egg-Based Production of Influenza Vaccine: 30 Years of Commercial Experience Cell-Culture-Based Vaccine Production: Technological Options Alternative Methods of Making Influenza Vaccines Adapting Industry Practices for the Rapid, Large-Scale Manufacture of Pharmaceutical Proteins,” 2006. [Online]. Available: http://www.nae.edu/TheBridge.

[10] M. M. J. Cox and J. R. Hollister, “FluBlok, a next generation influenza vaccine manufactured in insect cells,” Biologicals, vol. 37, no. 3, pp. 182–189, Jun. 2009, doi: 10.1016/J.BIOLOGICALS.2009.02.014.

[11] E. A. S. Nelson et al., “A pilot randomized study to assess immunogenicity, reactogenicity, safety and tolerability of two human papillomavirus vaccines administered intramuscularly and intradermally to females aged 18–26 years,” Vaccine, vol. 31, no. 34, pp. 3452–3460, Jul. 2013, doi: 10.1016/J.VACCINE.2013.06.034.

[12] D. V. Parums, “Editorial: First Approval of the Protein-Based Adjuvanted Nuvaxovid (NVX-CoV2373) Novavax Vaccine for SARS-CoV-2 Could Increase Vaccine Uptake and Provide Immune Protection from Viral Variants,” Med Sci Monit, vol. 28, pp. e936523–1, 2022, doi: 10.12659/MSM.936523.

[13] J. G. Choi et al., “Protective efficacy of baculovirus-derived influenza virus-like particles bearing H5 HA alone or in combination with M1 in chickens,” Vet Microbiol, vol. 162, no. 2–4, pp. 623–630, Mar. 2013, doi: 10.1016/J.VETMIC.2012.11.035.

[14] M. Subathra, P. Santhakumar, S. Satyam Naidu, M. Lakshmi Narasu, T. M. A. Senthilkumar, and S. K. Lal, “Expression of avian influenza virus (H5N1) hemagglutinin and matrix protein 1 in Pichia pastoris and evaluation of their immunogenicity in mice,” Appl Biochem Biotechnol, vol. 172, no. 7, pp. 3635–3645, Feb. 2014, doi: 10.1007/S12010-014-0771-Z/FIGURES/5.

[15] N. Sriwilaijaroen and Y. Suzuki, “Molecular basis of the structure and function of H1 hemagglutinin of influenza virus,” *Proceedings of the Japan Academy*, Series B, vol. 88, no. 6, pp. 226–249, Jun. 2012, doi: 10.2183/PJAB.88.226.

[16] L. V. Kordyukova, E. V. Shtykova, L. A. Baratova, D. I. Svergun, and O. V. Batishchev, “Matrix proteins of enveloped viruses: a case study of Influenza A virus M1 protein,” J Biomol Struct Dyn, vol. 37, no. 3, pp. 671–690, Feb. 2019, doi: 10.1080/07391102.2018.1436089.

[17] J.-M. Song et al., “Influenza Virus-Like Particles Containing M2 Induce Broadly Cross Protective Immunity,” PLoS One, vol. 6, no. 1, p. e14538, Jan. 2011, doi: 10.1371/journal.pone.0014538.

[18] E. Milián and A. A. Kamen, “Current and Emerging Cell Culture Manufacturing Technologies for Influenza Vaccines,” Biomed Res Int, vol. 2015, no. 1, p. 504831, Jan. 2015, doi: 10.1155/2015/504831.

[19] R. Correia et al., “Continuous Production of Influenza VLPs Using IC-BEVS and Multi-Stage Bioreactors,” Biotechnol Bioeng, vol. 122, no. 4, pp. 846–857, Apr. 2025, doi: 10.1002/BIT.28925;PAGEGROUP:STRING:PUBLICATION.

[20] O. Khanal and A. M. Lenhoff, “Developments and opportunities in continuous biopharmaceutical manufacturing.,” MAbs, vol. 13, no. 1, p. 1903664, 2021, doi: 10.1080/19420862.2021.1903664.

[21] L. Gerstweiler, J. Billakanti, J. Bi, and A. P. J. Middelberg, “An integrated and continuous downstream process for microbial virus-like particle vaccine biomanufacture.,” Biotechnol Bioeng, vol. 119, no. 8, pp. 2122–2133, Aug. 2022, doi: 10.1002/bit.28118.

[22] C. Matanguihan and P. Wu, “Upstream continuous processing: recent advances in production of biopharmaceuticals and challenges in manufacturing,” Curr Opin Biotechnol, vol. 78, p. 102828, Dec. 2022, doi: 10.1016/J.COPBIO.2022.102828.

[23] D. G. Turpeinen et al., “Continuous purification of an enveloped and non-enveloped viral particle using an aqueous two-phase system,” Sep Purif Technol, vol. 269, p. 118753, Aug. 2021, doi: 10.1016/J.SEPPUR.2021.118753.

[24] N. M. Nold et al., “Continuous purification of a parvovirus using two aqueous two-phase extraction steps,” Biotechnol Prog, vol. 41, no. 5, Sep. 2025, doi: 10.1002/btpr.70034.

[25] K. Aggarwal, F. Jing, L. Maranga, and J. Liu, “Bioprocess optimization for cell culture based influenza vaccine production,” Vaccine, vol. 29, no. 17, pp. 3320–3328, Apr. 2011, doi: 10.1016/j.vaccine.2011.01.081.

[26] F. Tapia, D. Wohlfarth, V. Sandig, I. Jordan, Y. Genzel, and U. Reichl, “Continuous influenza virus production in a tubular bioreactor system provides stable titers and avoids the ‘von Magnus effect,’” PLoS One, vol. 14, no. 11, p. e0224317, Nov. 2019, doi: 10.1371/journal.pone.0224317.

[27] C. C. Lai et al., “Process development for pandemic influenza VLP vaccine production using a baculovirus expression system,” J Biol Eng, vol. 13, no. 1, pp. 1–9, Oct. 2019, doi: 10.1186/S13036-019-0206-Z/TABLES/3.

[28] P. J. Krell, “Passage effect of virus infection in insect cells,” Cytotechnology, vol. 20, no. 1–3, pp. 125–137, 1996, doi: 10.1007/BF00350393/METRICS.

[29] P. von Magnus, “Incomplete Forms of Influenza Virus,” 1954, pp. 59–79. doi: 10.1016/S0065-3527(08)60529-1.

[30] A. Roldão, T. Vicente, C. Peixoto, M. J. T. Carrondo, and P. M. Alves, “Quality control and analytical methods for baculovirus-based products,” J Invertebr Pathol, vol. 107, no. SUPPL., pp. S94–S105, Jul. 2011, doi: 10.1016/J.JIP.2011.05.009.

[31] H. C. Wu, Y. C. Hu, and W. E. Bentley, “Tubular bioreactor for probing baculovirus infection and protein production,” Methods in Molecular Biology, vol. 1350, pp. 461–467, Jan. 2016, doi: 10.1007/978-1-4939-3043-2_23/FIGURES/2.

[32] F. Tapia, D. Wohlfarth, V. Sandig, I. Jordan, Y. Genzel, and U. Reichl, “Continuous influenza virus production in a tubular bioreactor system provides stable titers and avoids the ‘von Magnus effect,’” PLoS One, vol. 14, no. 11, p. e0224317, Nov. 2019, doi: 10.1371/journal.pone.0224317.

[33] Y.-C. Hu, M.-Y. Wang, and W. E. Bentley, “A tubular segmented-flow bioreactor for the infection of insect cells with recombinant baculovirus,” Cytotechnology, vol. 24, no. 2, pp. 143–152, 1997, doi: 10.1023/A:1007970020274.

[34] Life Technologies Corporation, “ExpiSfTM Expression System USER GUIDE,” 2018.

[35] W. Qingsheng and L. Yuanyuan, “Establishment, verification and application of rapid detection of baculovirus infectious titer by flow cytometry,” J Virol Methods, vol. 303, p. 114495, May 2022, doi: 10.1016/j.jviromet.2022.114495.

[36] P. J. Bottino, “Gus Gene Assay in Transformed Tissues,” https://goldbio.com/documents/1053/Gus%20Gene%20Assay%20Protocol.pdf.

[37] “ab234625 β-Glucuronidase Activity Assay Kit (Fluorometric),” https://www.abcam.com/en-us/products/assay-kits/glucuronidase-activity-assay-kit-fluorometric-ab234625.

[38] P. Ladiwala et al., “Unraveling Cytotoxicity in HEK293 Cells During Recombinant AAV Production for Gene Therapy Applications,” Biotechnol J, vol. 20, no. 3, Mar. 2025, doi: 10.1002/biot.202400501.

[39] S. Bio-Imaging, “Negative Staining,” http://web.path.ox.ac.uk/~bioimaging/bitm/instructions_and_information/EM/neg_stain.pdf.

[40] C. A. Schneider, W. S. Rasband, and K. W. Eliceiri, “NIH Image to ImageJ: 25 years of image analysis,” Nat Methods, vol. 9, no. 7, pp. 671–675, Jul. 2012, doi: 10.1038/nmeth.2089.

[41] H. Motulsky and A. Christopoulos, Fitting Models to Biological Data Using Linear and Nonlinear Regression. Oxford University PressNew York, NY, 2004. doi: 10.1093/oso/9780195171792.001.0001.

[42] O. Levenspiel, “Chemical reaction engineering / Octave Levenspiel.,” Chemical reaction engineering /, 2001, Accessed: Jul. 17, 2025. [Online]. Available: https://books.google.com/books/about/Chemical_Reaction_Engineering.html?id=vw48EAAAQBAJ

[43] A. E. Rodrigues, “Residence time distribution (RTD) revisited,” Chem Eng Sci, vol. 230, p. 116188, Feb. 2021, doi: 10.1016/j.ces.2020.116188.

[44] Y. Voloshin, A. Lawal, and N. S. Panikov, “Continuous plug-flow bioreactor: Experimental testing with Pseudomonas putida culture grown on benzoate,” Biotechnol Bioeng, vol. 91, no. 2, pp. 254–259, Jul. 2005, doi: 10.1002/BIT.20521;WGROUP:STRING:PUBLICATION.

[45] K. Trinh, M. Garcia-Briones, J. J. Chalmers, and F. Hink, “Quantification of damage to suspended insect cells as a result of bubble rupture,” Biotechnol Bioeng, vol. 43, no. 1, pp. 37–45, Jan. 1994, doi: 10.1002/bit.260430106.

[46] J. J. Chalmers and F. Bavarian, “Microscopic Visualization of Insect Cell-Bubble Interactions. II: The Bubble Film and Bubble Rupture,” Biotechnol Prog, vol. 7, no. 2, pp. 151–158, Mar. 1991, doi: 10.1021/bp00008a010.

[47] M. Muradoglu, A. Günther, and H. A. Stone, “A computational study of axial dispersion in segmented gas-liquid flow,” Physics of Fluids, vol. 19, no. 7, Jul. 2007, doi: 10.1063/1.2750295.

[48] D. L. Jarvis and A. Garcia, “Biosynthesis and Processing of the Autographa californica Nuclear Polyhedrosis Virus gp64 Protein,” Virology, vol. 205, no. 1, pp. 300–313, Nov. 1994, doi: 10.1006/viro.1994.1646.

[49] S. A. Monsma, A. G. Oomens, and G. W. Blissard, “The GP64 envelope fusion protein is an essential baculovirus protein required for cell-to-cell transmission of infection,” J Virol, vol. 70, no. 7, pp. 4607–4616, Jul. 1996, doi: 10.1128/jvi.70.7.4607-4616.1996.

[50] K. U. Dee and M. L. Shuler, “A mathematical model of the trafficking of acid-dependent enveloped viruses: Application to the binding, uptake, and nuclear accumulation of baculovirus,” Biotechnol Bioeng, vol. 54, no. 5, pp. 468–490, Jun. 1997, doi: 10.1002/(SICI)1097-0290(19970605)54:5<468::AID-BIT7>3.0.CO;2-C.

[51] L. K. Nielsen, “Virus Production from Cell Culture, Kinetics,” in *Encyclopedia of Cell Technology*, Wiley, 2000. doi: 10.1002/0471250570.spi106.

[52] Y. V. Lyupina, S. B. Abaturova, P. A. Erokhov, O. V. Orlova, S. N. Beljelarskaya, and V. S. Mikhailov, “Proteotoxic stress induced by Autographa californica nucleopolyhedrovirus infection of Spodoptera frugiperda Sf9 cells,” Virology, vol. 436, no. 1, pp. 49–58, Feb. 2013, doi: 10.1016/j.virol.2012.10.018.

[53] B. Sperker, T. E. Mürdter, J. T. Backman, P. Fritz, and H. K. Kroemer, “Expression of active human β-glucuronidase in Sf9 cells infected with recombinant baculovirus,” Life Sci, vol. 71, no. 13, pp. 1547–1557, Aug. 2002, doi: 10.1016/S0024-3205(02)01917-3.

[54] T. K. K. Wong, L. K. Nielsen, P. F. Greenfield, and S. Reid, “Relationship between oxygen uptake rate and time of infection of Sf9 insect cells infected with a recombinant baculovirus,” Cytotechnology, vol. 15, no. 1–3, pp. 157–167, 1994, doi: 10.1007/BF00762390.

[55] X. Li, J. Gu, H. Wu, and Y. Xie, “Pilot-scale process development for recombinant adeno-associated virus (rAAV) production based on high-density Sf9 cell culture,” Virol J, vol. 21, no. 1, p. 281, Nov. 2024, doi: 10.1186/s12985-024-02550-4.

[56] T. Gotoh, M. Fukuhara, and K. I. Kikuchi, “Mathematical model for change in diameter distribution of baculovirus-infected Sf-9 insect cells,” Biochem Eng J, vol. 40, no. 2, pp. 379–386, Jun. 2008, doi: 10.1016/J.BEJ.2008.01.008.

[57] R. D. Possee, A. C. Chambers, L. P. Graves, M. Aksular, and L. A. King, “Recent Developments in the Use of Baculovirus Expression Vectors,” Curr Issues Mol Biol, pp. 215–230, 2020, doi: 10.21775/cimb.034.215.

[58] B. J. Lindsay et al., “Morphological characterization of a plant-made virus-like particle vaccine bearing influenza virus hemagglutinins by electron microscopy,” Vaccine, vol. 36, no. 16, pp. 2147–2154, Apr. 2018, doi: 10.1016/J.VACCINE.2018.02.106.

[59] J. F. Power, S. Reid, K. M. Radford, P. F. Greenfield, and L. K. Nielsen, “Modeling and optimization of the baculovirus expression vector system in batch suspension culture,” Biotechnol Bioeng, vol. 44, no. 6, pp. 710–719, Sep. 1994, doi: 10.1002/bit.260440607.

[60] F. Destro et al., “Mechanistic modeling explains the production dynamics of recombinant adeno-associated virus with the baculovirus expression vector system.,” Mol Ther Methods Clin Dev, vol. 30, pp. 122–146, Sep. 2023, doi: 10.1016/j.omtm.2023.05.019.

[61] S. Kiesslich and A. A. Kamen, “Vero cell upstream bioprocess development for the production of viral vectors and vaccines,” Biotechnol Adv, vol. 44, p. 107608, Nov. 2020, doi: 10.1016/j.biotechadv.2020.107608.

[62] J. Ou et al., “Scalable process development for rAAV transient transfection production using computational fluid dynamics modeling,” Biotechnol Prog, vol. 41, no. 5, Sep. 2025, doi: 10.1002/btpr.70028.

[63] O. F. Garcia-Aponte, C. Herwig, and B. Kozma, “Lymphocyte expansion in bioreactors: upgrading adoptive cell therapy,” J Biol Eng, vol. 15, no. 1, p. 13, Apr. 2021, doi: 10.1186/s13036-021-00264-7.

[64] G. Nadal-Rey, D. D. McClure, J. M. Kavanagh, S. Cornelissen, D. F. Fletcher, and K. V. Gernaey, “Understanding gradients in industrial bioreactors,” Biotechnol Adv, vol. 46, p. 107660, Jan. 2021, doi: 10.1016/j.biotechadv.2020.107660.

[65] V. Girard et al., “Large-scale monoclonal antibody purification by continuous chromatography, from process design to scale-up,” J Biotechnol, vol. 213, pp. 65–73, Nov. 2015, doi: 10.1016/j.jbiotec.2015.04.026.

